# Semantic Representation of Neural Circuit Knowledge in *Caenorhabditis elegans*

**DOI:** 10.1101/2023.04.28.538760

**Authors:** Sharan J. Prakash, Kimberly M. Van Auken, David P. Hill, Paul W. Sternberg

## Abstract

In modern biology, new knowledge is generated quickly, making it challenging for researchers to efficiently acquire and synthesise new information from the large volume of primary publications. To address this problem, computational approaches that generate machine-readable representations of scientific findings in the form of knowledge graphs have been developed. These representations can integrate different types of experimental data from multiple papers and biological knowledge bases in a unifying data model, providing a complementary method to manual review for interacting with published knowledge. The Gene Ontology Consortium (GOC) has created a semantic modelling framework that extends individual functional gene annotations to structured descriptions of causal networks representing biological processes (Gene Ontology Causal Activity Modelling, or GO-CAM). In this study, we explored whether the GO-CAM framework could represent knowledge of the causal relationships between environmental inputs, neural circuits and behavior in the model nematode *C. elegans* (*C. elegans* Neural Circuit Causal Activity Modelling (*Ce*N- CAM)). We found that, given extensions to several relevant ontologies, a wide variety of author statements from the literature about the neural circuit basis of egg-laying and carbon dioxide (CO_2_) avoidance behaviors could be faithfully represented with *Ce*N-CAM. Through this process, we were able to generate generic data models for several categories of experimental results. We also discuss how semantic modelling may be used to functionally annotate the *C. elegans* connectome. Thus, Gene Ontology-based semantic modelling has the potential to support various machine-readable representations of neurobiological knowledge.

**Graphical Abstract:** 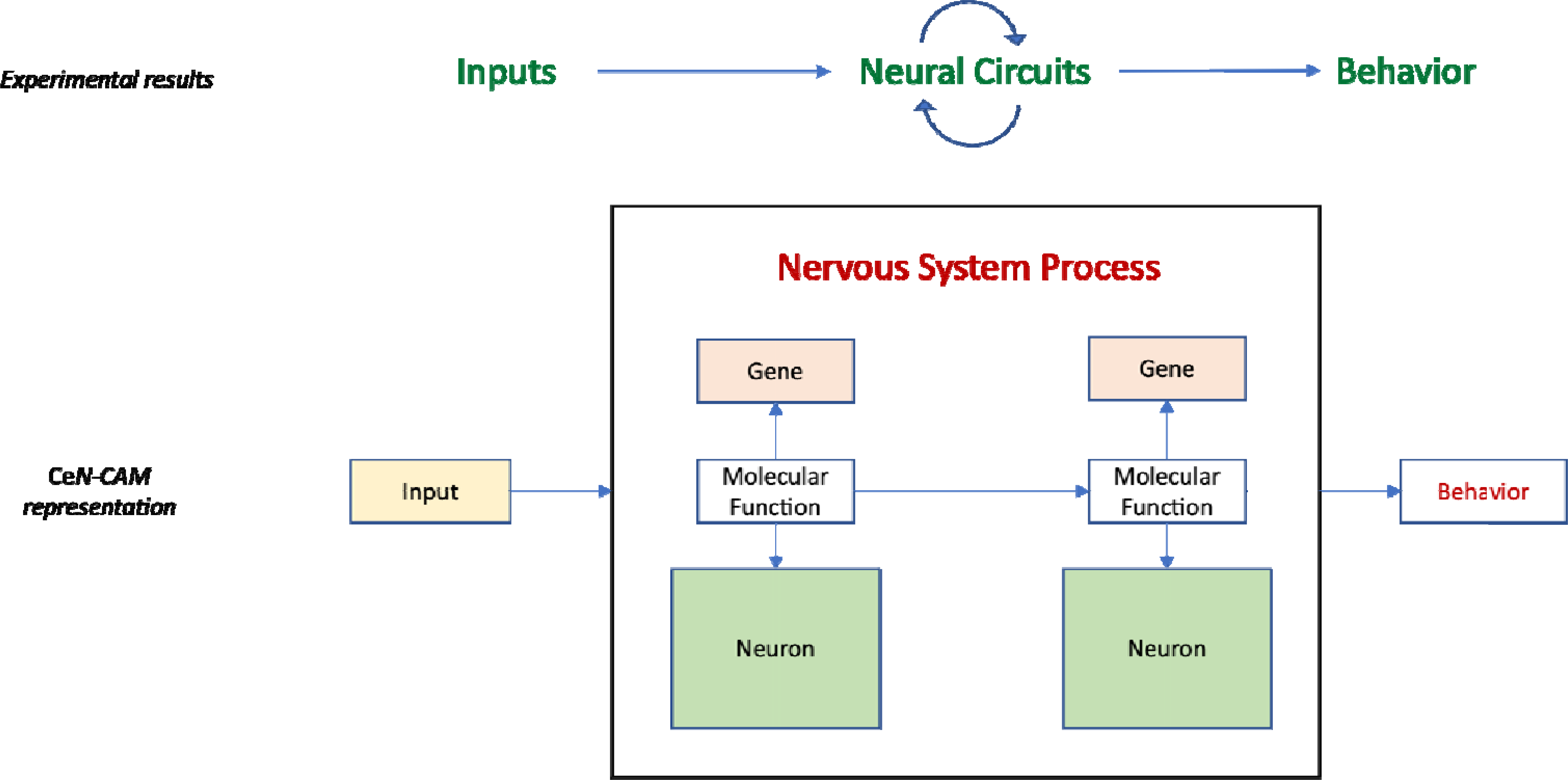

## Introduction

### *Caenorhabditis elegans* as a Model for Systems Neuroscience

A major goal of modern neuroscience is to explain the relationship between environmental inputs and complex behaviors in terms of the properties of their underlying neural systems. *C. elegans* has been a productive model for neuroscience due to its wide range of easily measured behaviors, genetic tractability, and highly stereotyped anatomy. The function of individual *C. elegans* neurons has been studied by a variety of methods, including selective neuron ablation, either with laser microbeam irradiation (Chalfie *et al*. 1985; Bargmann and Horvitz 1991; Bargmann and Avery 1995; Liu and Sternberg 1995) or genetically encoded cell killing (Harbinder et al. 1997; Srinivasan et al. 2012). These physical studies, complemented by genetic screens resulting in mutant animals with distinct behavioral and neuronal phenotypes, have implicated specific neurons in behaviors (Bargmann 1993) and identified genes and neurons required for responses to environmental or pharmacological inputs (Waggoner *et al*. 1998). Technological advances, such as cell-specific application of optogenetic and chemical perturbations (Husson *et al*. 2013; Pokala *et al*. 2014) in combination with calcium imaging of individual neurons (Chung *et al*. 2013), have begun to outline the causal relationships between neurons, both locally and via long-range connections (Shen *et al*. 2016), while calcium imaging allows the effect of physical inputs on neural activity to be determined. Thus, causal relationships can be traced from inputs through neural circuits to behavior. In addition, traditional molecular genetic methods enable the biochemical basis of these causal relationships to be elucidated. Understanding molecular participants is particularly important for the functional description of extra-synaptic connections because they cannot be described by anatomy or gene expression alone, yet they exert powerful effects on neuronal activity (Bargmann 2012; Marder 2012; Bentley *et al*. 2016). In combination, the physical and molecular data allow detailed description of *C. elegans* neural circuits underlying particular behaviors.

### The GO-CAM framework can be used to Represent Causal Relationships in Biology

Given the volume of biological knowledge, a method to integrate diverse types of data into causal models of biological systems, expressed in a common, machine-readable language, is highly desirable. A promising method suitable for this application has been developed. The Gene Ontology (GO) Consortium has created a semantic modelling framework for annotating causal relationships between molecular activities in the context of functional gene annotation, known as GO-CAM (Gene Ontology Causal Activity Modelling) (Thomas *et al*. 2019).

Semantic models (also known as knowledge graphs) are machine-readable representations of knowledge in a given field, in which the edges of the graph describe the logical relationships between entities that comprise a field of study. In GO-CAM, curated knowledge of gene functions annotated using the Gene Ontology and other biologically relevant ontologies are used to create activity flow models of biological systems (Fig. 1) (Le Novère et al. 2009). In these graphs, the logical relationships are described via a formalism known as a semantic triple (subject-predicate-object^1^). These models can be thought of as compositions of assertions in the form of semantic triples. For instance, the assertion “[*G-protein coupled receptor activity* (GO:0004930)] *has input* [*2-heptanone* (CHEBI: 5672)]” is a semantic triple that could be included in a GO-CAM. The semantic triple format allows edges to connect many different kinds of entities, including anatomy terms and biological processes. For instance, “[*glucose-6-phosphate isomerase activity* (GO:0004374)] *part of* [*canonical glycolysis* (GO:0061621)] *occurs in* [*cytosol* (GO:0005829)]” is a pair of semantic triples that connects a GO molecular function to both a higher-level biological process and an anatomical compartment. The Gene Ontology itself follows a hierarchical structure described with semantic triples, e.g. “[*G-protein coupled receptor activity* (GO:0004930)] *is a* [*transmembrane signalling receptor* (GO:0004888)]” (here the relation ‘*is a*’ describes a child-parent relationship). This formalism allows different kinds of entities to be connected to one another in a machine-readable format, allowing combinatorial queries and other computational analyses.

**Figure 1.**
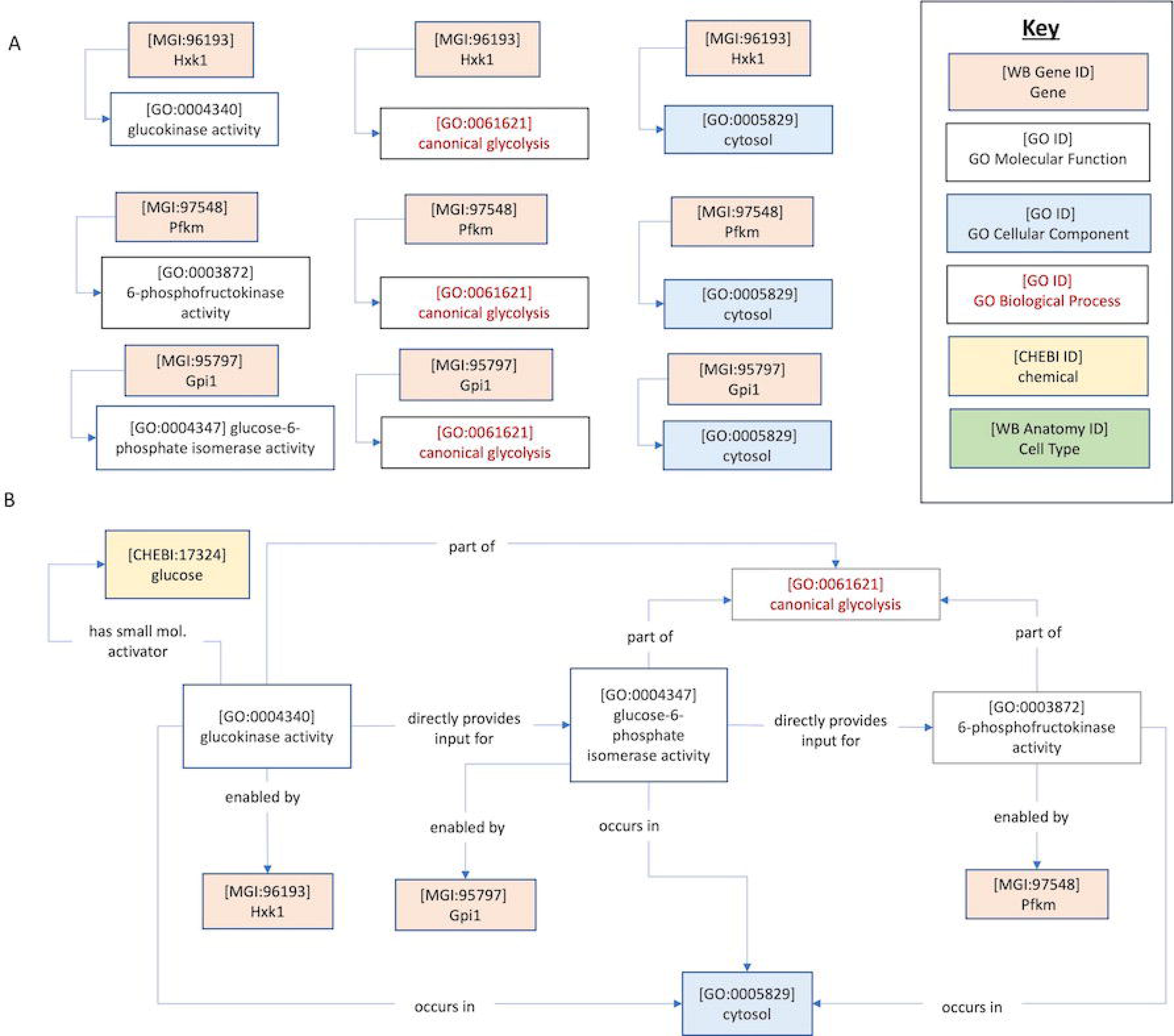
Standard GO annotations and GO-CAMs A) Standard GO annotations link genes to GO Molecular Functions, GO Biological Processes or GO Anatomy terms. B) A partial GO-CAM of canonical glycolysis. Gene Ontology-Causal Activity Models (GO-CAMs) arrange GO annotations into structured models of biological processes by causally linking GO molecular functions that make up a process. Edges represent relations which may connect nodes according to the GO-CAM data model.

In GO-CAM, curated knowledge of gene functions annotated using the Gene Ontology and other biologically relevant ontologies are used to create knowledge graphs of biological systems (Fig. 1). This framework extends traditional gene function annotation by capturing the causal flow of molecular activities, e.g. protein kinase activity or ion channel activity, using causal relations from the Relations Ontology (RO) and representing these interactions in the context of the relevant biological process and anatomy (Smith *et al*. 2005) (Box 1). These causal networks allow more in-depth computational analyses of a system than a set of stand-alone associations between genes and ontology terms, and have the potential to bridge the gap between biochemical and anatomical networks. Here, we explored whether the causal GO-CAM framework can enable the representation of the causal relationships between environmental inputs, neural circuits and behavior at varying levels of detail.

#### Box 1.

**Commonly used RO Relations in GO-CAM**

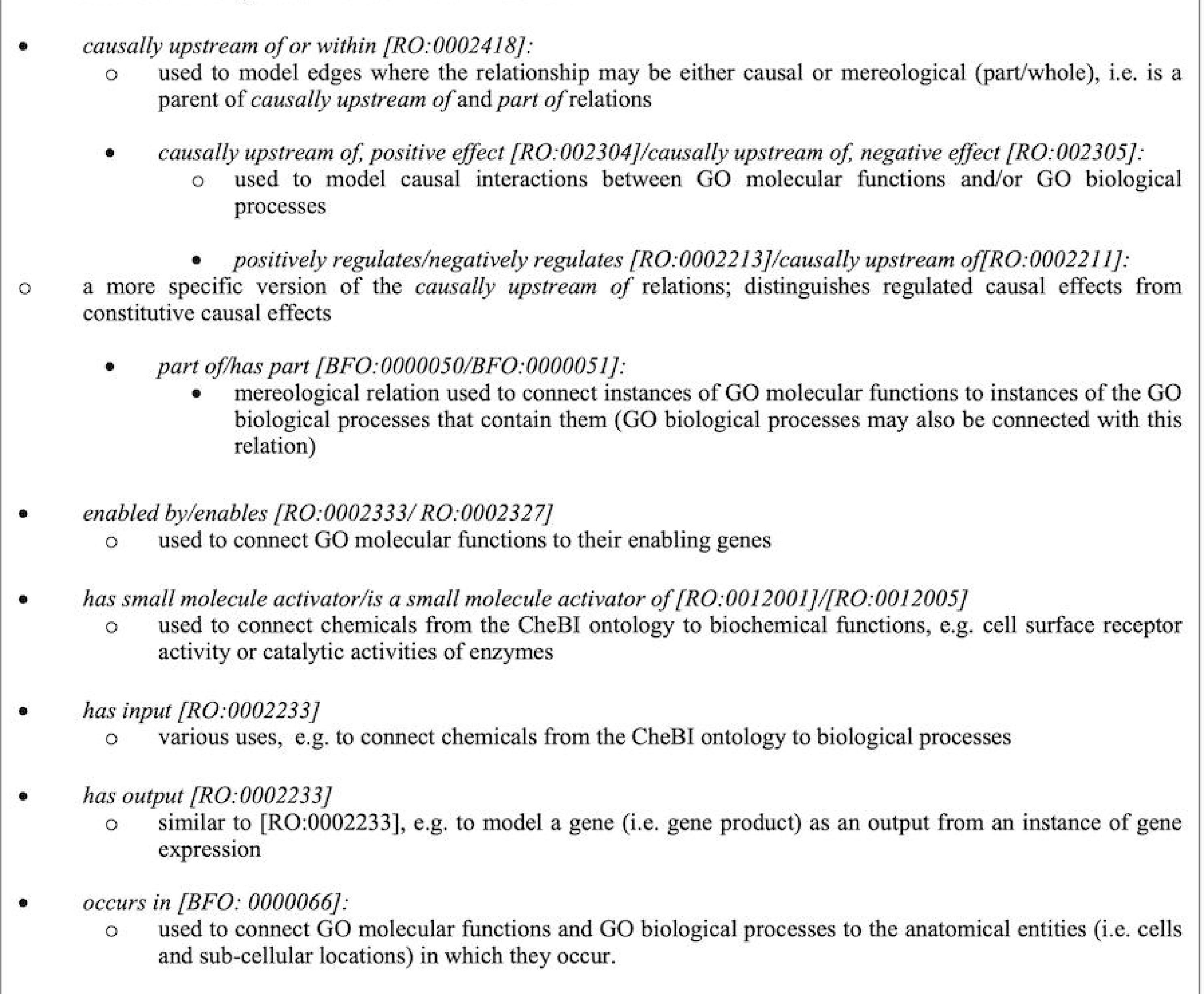

### *Ce*N-CAM: GO-CAM Representation of *C. elegans* Neurobiological Knowledge

As for standard GO annotations, assertions in a GO-CAM are supported by evidence statements, ideally experimental evidence from the published literature (Ashburner *et al*. 2000; Giglio *et al*. 2019; The Gene Ontology Consortium 2021). To adapt the GO-CAM framework for modelling neurobiological statements about *C. elegans* egg-laying and carbon dioxide (CO_2_)-sensing behaviors, we selected a subset of relevant papers from the *C. elegans* bibliography and identified author statements that could be used to support construction of semantically rigorous, causal models. For the egg-laying circuit, these statements largely involve interactions among interneurons, motor neurons, and the egg-laying apparatus, e.g. vulval muscles and epithelia. The CO_2_ avoidance circuit is focused on sensory neurons, their interaction with the environment, and subsequent effects on locomotory behavior.

## Methods & Materials

To model neurobiological processes, we began by collecting author statements from published references. In order to ensure that our findings were broadly applicable, we collected statements from the literature on two circuits, one centred on interneurons and motor neurons (egg-laying) and one centred on sensory neurons (CO_2_ avoidance). For the egg-laying circuit we compiled 20 papers, and for CO_2_ avoidance, 8 papers. We chose statements manually, according to a few criteria. To begin with, we chose statements that provided a clear interpretation and that we therefore expected to be straightforward to model with GO-CAM. Later, we selected statements describing phenomena (e.g. multi-sensory integration, neuromodulation) that were missing from the initial dataset.

We defined an author statement as text describing: i) either an experiment or hypotheses, ii) an experimental observation or result, and iii) a clear biological interpretation of the result. These typically comprised a paragraph. We then attempted to model the interpretation, along with supporting evidence using the Evidence and Conclusion Ontology (ECO) (Giglio *et al*. 2019) wherever possible. We avoided modelling speculative suggestions that went beyond the supporting evidence.

For each author statement, we attempted to generate one or more simple assertions (i.e. semantic triples or subject-predicate-object) that accurately modelled the author statement using classes from biological ontologies (Table 1) including the GO (Ashburner *et al*. 2000; The Gene Ontology Consortium 2021), the Chemicals of Biological Interest ontology (ChEBI) (Hastings et al. 2016), the Environmental Conditions, Treatments & Exposures Ontology (ECTO) (Chan *et al*. 2022), and the *C. elegans* Cell and Anatomy Ontology (WBbt) (Lee and Sternberg 2003). In a semantic triple, these classes are connected by relations from the Relations Ontology (Smith *et al*. 2005) (Box 1).

**Table 1:** Biological Ontologies Used To Generate CeN-CAM models.

| GO-CAM element | Ontology | Example |
| --- | --- | --- |
| Molecular activity | GO molecular function | serotonin receptor activity (GO:0099589) |
| Biological process | GO biological process | Membrane depolarization (GO:0051899) |
| Location | GO cellular component | cytosol (GO:0005829) |
| Cell | WormBase Anatomy Ontology | HSN (WBbt:0006830) |
| Active Entity (Gene/Gene Product) | WormBase | *tph-1* (WBGene00006600) |
| Chemical inputs | Chemical Entities of Biological Interest (ChEBI) | dioxygen (ChEBI:15379) |
| Relations arrows | Relations Ontology (RO), Basic Formal Ontology (BFO) | occurs in (RO_0002479) |
| Evidence codes | Evidence & Conclusions Ontology (ECO) | optogenetic evidence used in manual assertion (ECO:0006033) |
| Environmental Conditions | Phenotype and Trait Ontology (PATO)<br>Environmental Conditions, Treatments and Exposure Ontology (ECTO) | Increased duration (PATO_0000498) |

We collected author statements and their corresponding semantic triples into a dataframe such that the triple representation can be read from left to right (unless otherwise specified). Supplementary Tables 1 and 2 provide the full list of author statements that were modelled for the egg-laying (91 unique statements comprising 128 entries from 20 papers) and CO_2_-avoidance circuits (59 unique statements comprising 99 entries from 8 papers), respectively. Table 2 enumerates detailed categories of biological phenomena captured by this approach. We used this categorization process to determine whether existing ontologies contained a sufficiently rich set of classes and whether existing RO terms were adequate to describe the relations between classes. Where applicable, we generated definitions for required novel classes and their necessary parents (Table 3). We then created illustrations of several useful examples.

**Table 2:** Categories of Neurobiological Phenomena Modelled with GO-CAM.

|  |  |
| --- | --- |
| **Neuronal basis of behavior** | **Receptor or G-protein activity regulates behavior or cell activity** |
| Neuron regulates behavior | G protein activity in specific neuron regulates behavior |
| Cellular process regulates behavior | GPCR regulates G -protein-activity in specific neuron to regulate behavior |
| Neuronal activity regulates behavior | Neuromodulation of specific neuron by G protein signaling |
| Neuronal activity dependent secretion from identified neuron regulates behavior | G protein activity regulates gene expression |
| Neuronal activity dependent neurotransmitter secretion | G protein activity regulates gene expression cell autonomously |
| **Neuron-neuron interaction** | G protein activity regulates neurotransmitter biosynthesis |
| Activity of Neuron A regulates activity of Neuron B | G protein activity regulates phospholipase activation |
| Activity of Neuron A regulates activity of Neuron B (synapse-dependent) | G protein signaling activity regulates neuronal activity cell autonomously |
| Mechanical stimulation of Neuron A regulates activity of Neuron B | GPCR regulates ion channel |
| Negative autoregulation of neuronal activity | Ion channel regulates neuronal activity via GPCR |
| Neural activity depends on extra-synaptic signaling | G protein activity regulates neurotransmitter biosynthesis cell autonomously |
| **Environmental influence on behavior or cell activity** | GPCR regulates G -protein-activity |
| Environmental input regulates behavior | G protein activity regulates neurotransmitter biosynthesis |
| Environmental input regulates neuronal activity | Neuromodulation of specific neuron by G protein signaling |
| Environmental condition regulates neuronal activity | Receptor activity regulates neuronal activity cell autonomously |
| Environmental condition regulates behavior | **Cellular Process** |
| Mechanical process regulates neural activity | Neurotransmitter biosynthesis |
| Environmental input regulates behavior via defined neuron | Neurotransmitter signaling pathway affects behavior |
| Environmental input regulates gene expression | Biochemical process regulates neural activity |
| **Receptor-ligand Interaction** | Dense core vesicle exocytosis from identified neuron regulates behavior |
| Receptor-ligand interaction | Gene activity regulates neural activity |
| Neurotransmitter regulates neuronal activity via ion channel | Gene activity in identified neuron regulates behavior |
| Neurotransmitter regulates behavior via specific receptor | Dense core vesicle exocytosis regulates behavior |
| Neurotransmitter regulates behavior via ion channel in identified neuron | Neuropeptide signaling pathway affects behavior |
| Neurotransmitter regulates behavior via specific receptor in identified neuron | Neuropeptide signaling pathway affects behavior via identified cell |
| Neurotransmitter affects identified receptor class | Regulation of gene expression in identified neuron |
| **Ion channel activity regulates behavior or cell activity** | **Neurotransmitter/neuropeptide activity regulates behavior or cell activity** |
| Ion channel regulates neural activity | Neurotransmitter biosynthesis from identified source neuron regulates behavior |
| Neuromodulation of specific neuron by ion channel activity | Neurotransmitter biosynthesis regulates behavior |
| Ion channel regulates membrane potential | Neuropeptide from specific neuron regulates behavior |
| Ion channel activity in defined neuron regulates behavior | Neurotransmitter activity depends on ion channel |
| Ion channel activity regulates behavior | Neurotransmitter regulates behavior |
| **Nervous System Process** | Neurotransmitter regulates neuronal activity |
| Adaptation to chemical stimulus | Regulation of neurotransmitter activity by upstream neuropeptide activity |
| Co-ordination of locomotion and neural activity to influence behavior | Regulation of secretion by upstream neuropeptide activity |

**Table 3:** Definitions & Classification for Proposed New GO Classes.

| Category | Proposed Modified Classes | Modification | Classification |
| --- | --- | --- | --- |
| GO Biological Process | *Oviposition (GO:0018991)* | Change primary term name to synonym 'egg-laying behavior' New definition: The muscle system process resulting in the deposition of eggs (either fertilized or not) upon a surface or into a medium such as water. | is a *muscle system process (GO:0003012)*, part of 'egg-laying behavior' |
| GO Biological Process | *Behavior (GO:0007610)* | New classification (see right) | is a *response to stimulus (GO:0050896)* |
| Category | Proposed New Class | Definitions | Classification |
| GO Biological Process | Egg deposition | The multicellular organismal reproductive process that results in the movement of an egg from within an organism into the external environment. | is a *reproductive behavior (GO:00198098)*; part of *oviposition/egg-laying behavior (GO:0018991)* |
| GO Biological Process | Positive regulation of egg deposition | Any process that positively regulates the rate, frequency or extent of egg deposition | is a *nervous system process* (GO:0050877); part of *oviposition/egg-laying behavior* (GO:0018991) |
| GO Biological Process | Negative regulation of egg deposition | Any process that negatively regulates the rate, frequency or extent of egg deposition | is a *nervous system process* (GO:0050877); part of *oviposition/egg-laying behavior* (GO:0018991) |
| GO Biological Process | Neuron-to-neuron extra-synaptic peptide signaling | Any process by which a cellular process within one neuron influences a cellular process within another neuron via a secreted gene product, where this secretion occurs independently of synapses | is a *neuron-to-neuron chemical signaling process* |
| GO Biological Process | Neuron-to-neuron chemical signaling | Any process by which a cellular process within one neuron influences a cellular process within another neuron via a secreted molecule | is a *cell-cell signaling* (GO:0007267) process |
| GO Biological Process | Behavioral response to carbon dioxide | The behavior of an organism in response to a carbon dioxide stimulus. | is a *behavior* (GO:007610); is a *response to carbon dioxide* (GO:0010037) |
| GO Biological Process | Carbon dioxide avoidance behavior | The behavioral response to carbon dioxide which results in the directed movement of a motile cell or organism towards a lower carbon dioxide concentration | is a *negative chemotaxis* (GO:0050919); is a *behavioral response to carbon dioxide* |
| GO Biological Process | Negative regulation of carbon dioxide avoidance behavior | Any process that negatively influences locomotory behavior directed away from a source or gradient of carbon dioxide . | is a *nervous system process* (GO:0050877); part of *behavioral response to carbon dioxide* |
| GO Biological Process | Positive regulation of carbon dioxide avoidance behavior | Any process that positively influences locomotory behavior directed away from a source or gradient of carbon dioxide. | is a *nervous system process* (GO:0050877); part of *behavioral response to carbon dioxide* |
| GO Biological Process | Adaptation of neuron to stimulus | Any process that results in an increased threshold for induction of neural activity due to prior exposure to the same stimulus | is a *negative regulation of membrane depolarization* (GO1904180) |
| GO Biological Process | Sensitization of neuronal response to stimulus | Any process that results in a reduced threshold for induction of neural activity due to prior exposure to the same stimulus. | is a *positive regulation of response to stimulus* (GO:0048584) |
| GO Biological Process | Sensitization of behavioral response to stimulus | Any process that results in a reduced threshold for induction of behavioral response due to prior exposure to the same stimulus. | is a *positive regulation of behavior* (GO:0048520) |
| GO Biological Process | Sensory adaptation in behavioral response to stimulus | Any process that results in an increased threshold for induction of behavioral response due to prior exposure to the same stimulus. | is a *negative regulation of behavior* (GO:0048521) |
| GO Biological Process | Behavior co-ordination process | Any neural process that links the execution or cessation of one behavior to the induction or cessation of another behavior. | is a *regulation of behavior* (GO:0050795) |
| GO Biological Process | Signal integration Process | Any nervous system process by which different types of input to the nervous system contribute in combination to a behavioral or physiological output. | is a *nervous system process* (GO:0050877) |
| GO Molecular Function | CO2 receptor activity | Binding to and responding, e.g. by conformational change, to changes in the cellular level of carbon dioxide (CO2) or its dissociation products in water. | is a *signaling receptor activity* (GO:0038023) |
| ECO Evidence Class | Neuron Chemical Inhibition Assay Evidence used in Manual Assertion | A type of experimental phenotypic evidence that is used in a manual assertion, arising from experiments in which the output from a neuron is inhibited by a chemical | is a *experimental phenotypic evidence* (ECO:0000059) |
| ECO Evidence Class | Synaptic Transmission Inhibition Evidence used in Manual Assertion | A type of experimental phenotypic evidence, that is used in a manual assertion, arising from experiment in which neuron-to-neuron synaptic transmission is manipulated using inhibitors of synaptic transmission, that is used in a manual assertion. | is a *experimental phenotypic evidence* (ECO:0000059) |
| ECO Evidence Class | Mechanical Perturbation Evidence used in Manual Assertion | A type of experimental phenotypic evidence, that is used in a manual assertion, arising from experiment in which cellular responses are manipulated using mechanical force, , that is used in a manual assertion. | is a *experimental phenotypic evidence* (ECO:0000059) |
| ECO Evidence Class | Long-term Exposure or Conditioning Evidence used in Manual Assertion | A type of experimental phenotypic evidence, that is used in a manual assertion, arising from experimental treatment involving sustained exposure of an organism to one or more environmental conditions. | is a *experimental phenotypic evidence* (ECO:0000059) |
| ECTO Class | exposure to increasing carbon dioxide | An exposure event involving the interaction of an exposure receptor to increasing amount of carbon dioxide | is a *exposure to chemical* (ECTO: 0000231) |
| ECTO Class | exposure to decreasing carbon dioxide | An exposure event involving the interaction of an exposure receptor to increasing amount of carbon dioxide | is a *exposure to chemical* (ECTO: 0000231) |

In generating our empirical models, we sought as far as possible to ensure that all relations followed the conventions of the GO-CAM data model. Namely, two GO Molecular Functions can be linked by causal relations, whereas a GO Molecular Function (MF) and a GO Biological Process (BP) are linked by mereological relations (e.g. *part of*). In addition, two BPs can be linked by mereological relations when one BPs is part of another BP (i.e. a subprocess of the other). We also found it necessary, in some cases, to link distinct BPs using causal relations to accurately describe the complexity of the biology. For instance, one neuron activating another via optogenetics can be modelled by a *membrane depolarization* process causally upstream of another *membrane depolarization* process (e.g Fig. 4A). We sought to include whichever MFs or BPs were implied by an author statement, even if the gene was missing, or the BP was not explicitly discussed, in order to denote missing information. We chose the most specific relation or GO term that we felt was justified in the circumstances. For instance, when modelling individual author statements, we used *causally upstream of,* but when modelling compilations of statements from separate papers, we were able to use the child term *positively regulates*. In generating our generic template models, we chose the highest level relations and GO terms that could reasonably represent a given statement category.

**Table 4:** Author Statements Collection A.

|  | Author | Author Statement |
| --- | --- | --- |
| A | Waggoner *et al.* 1998 | "The roles of individual neurons in controlling the timing of egg-laying events can be determined with high precision by eliminating specific neurons by laser ablation and assaying the effect of the ablation on behavior. We therefore eliminated the neurons with prominent synaptic input to the egg-laying muscles to determine how their absence affected the timing of egg-laying events. We first investigated the involvement of the HSNs, a pair of serotonergic motor neurons that are required for efficient egg laying. By tracking the behavior of animals lacking both HSNs, we found that elimination of the HSNs did not qualitatively alter the pattern of egg laying: eggs were still laid in clusters, and the intervals between clusters and between egg-laying events within a cluster were still exponentially distributed. However, HSN ablation did cause a substantial lengthening of the inactive phase, which led to a slower overall rate of egg laying (Figure 2A). Since loss of the HSNs decreased the frequency of egg-laying clusters (i.e., $\lambda_2$ was decreased; Table 1) but did not slow the egg-laying rate within these clusters ($\lambda_1$ was actually increased), these results suggest that the HSNs stimulate egg laying by inducing the active state." |
| B | Bany *et al.* 2003 | "Because the VC neurons appear to inhibit egg laying and are cholinergic, we tested whether the VCs release acetylcholine to inhibit egg laying. The VCs are the only cells of the egg-laying system that express the UNC-4 complex, CHA-1, and UNC-17 (Lickteig *et al.*, 2001); however, because unc-4, cha-1, and unc-17 are each expressed in other neurons, it was necessary to determine whether mutations in these genes cause hyperactive egg laying specifically attributable to their effects on the VC neurons. For this purpose, we expressed the unc-4, cha-1, or unc-17 cDNAs in the VC neurons and determined whether this rescued the hyperactive egg-laying defects of the corresponding mutants. To direct VC expression, we used a modified lin-11 promoter similar to that used to express GFP in Figure 3A (see Materials and Methods). Expression of the unc-4 cDNA using this promoter rescued the hyperactive egg-laying defect of unc-4 mutants, returning the percentage of early-stage eggs laid to near-wild-type levels (Fig. 4A). Furthermore, expressing the cha-1 cDNA in the VC neurons of cha-1 mutants also rescued their hyperactive egg-laying phenotype (Fig. 4B). Similar experiments with unc-17 gave analogous results (data not shown). Restoring the inhibition of egg laying by restoring the ability of the VC neurons to signal with acetylcholine provides our most compelling evidence that it is the VC neurons that inhibit egg laying." |
| C | Banerjee *et al.* 2017 | "We next sought to determine whether uv1 activation is sufficient to inhibit egg-laying. To address this question, we expressed channelrhodopsin (ChR2) in uv1 cells using the regulatory regions of ocr-2 as above. Light stimulation immediately following the initial egg-laying event of an active phase (see Methods) significantly delays subsequent egg-laying events, and also significantly reduces the total number of egg-laying events within an active phase (Fig 2)(S1 Movie). For example, under control conditions a majority (~80%) of animals show a delay between the first and second egg-laying events within an active phase of <20 s (light stimulation, -ATR) (Fig 2B). This proportion is reduced dramatically (to around 10%) when uv1 cells are activated (light stimulation, +ATR)...Taken together, our findings provide evidence that uv1-mediated inhibition of egg-laying promotes periods of quiescence in the egg-laying program and plays a key role in setting their duration." |
| D | Carnell *et al.* 2005 | "The expression of gfp in the vulval muscles suggests that ser-1 may be acting in vulval muscles to mediate the stimulatory effect of 5-HT on egg laying. To test this hypothesis, we expressed the ser-1 cDNA using the muscle-specific myo-3 promoter (Okkema *et al.*, 1993) to determine whether it could rescue 5-HT-induced egg laying. Consistent with this hypothesis, we found the Pmyo-3::ser-1(+) transgene partially restored 5-HT-dependent egg laying to ser-1(ok345) animals (Fig. 2A). A wild-type ser-1 transgene with the same 3.4 kB promoter that failed to express gfp in the vulval muscles also failed to rescue the egg-laying defects of the ser-1 mutant animals (Fig. 2A). These results indicate that ser-1 expression in muscle can restore egg laying. Previous studies have indicated that 5-HT acts on vulval muscle to stimulate egg laying (Trent *et al.*, 1983; Brundage *et al.*, 1996; Waggoner *et al.*, 1998; Bastiani *et al.*, 2003; Shyn *et al.*, 2003). Our results indicate that ser-1 mediates this response." |
| E | Carillo *et al.* 2013 | "NPR-1 is not expressed in BAG neurons but is expressed in a number of other sensory neurons as well as some interneurons (Macosko *et al.*, 2009). To identify the site of action for the regulation of CO2 response by npr-1, we introduced the N2 allele of npr-1 into npr-1(lf) mutants in different subsets of neurons and assayed CO2 response. We found that expressing npr-1 in neuronal subsets that included the O2-sensing URX neurons (Cheung *et al.*, 2004; Gray *et al.*, 2004) restored CO2 response (Fig. 3A). These results suggest that NPR-1 activity in URX neurons is sufficient to enable CO2 avoidance. However, we cannot exclude the possibility that NPR-1 function in other neurons also contributes to CO2 avoidance." |
| F | Bretscher *et al.* 2011 | "Strikingly, AFD also responded to removal of CO2 with a fast Ca2+ spike that peaked within 10 s ("CO2-OFF" response)...CO2-evoked activity in AFD could be due to synaptic input to AFD. To test this, we imaged CO2 responses in unc-13 mutants, which have severe defects in synaptic release (Richmond *et al.*, 1999). The AFD CO2 responses of unc-13 animals were indistinguishable from wild-type (Figures 2H and S1C). These data suggest that, as well as being a thermosensory neuron (Mori and Ohshima, 1995; Kimura *et al.*, 2004; Clark *et al.*, 2007), AFD is a CO2 sensor with both ON and OFF responses." |
| G | Collins *et al.* 2016 | "To directly test how neurotransmitter signaling from the HSNs regulates egg-laying circuit activity, we used the egl-6 promoter to express Channelrhodopsin-2 in the HSNs (Emtage *et al.*, 2012), allowing us to drive neurotransmitter release specifically from the HSNs with blue light. ...We found that activation of HSNs resulted in circuit activity reminiscent of a spontaneous active state, including rhythmic Ca2+ activity of both VCs and vulval muscles, and egg-laying events that accompanied a subset of these Ca2+ transients... These results suggest that the high level of HSN activity after optogenetic |
|  |  | activation induces strong coupling of VC and vulval muscle excitation." |
| **H** | Kopchok *et al.* 2021 | "To determine whether VC synaptic transmission regulates egg laying via HSN, we recorded HSN Ca<sup>2+</sup> activity in WT and transgenic animals expressing TeTx in the VCs (Fig. 6A). During the egg-laying active state, the HSNs drive egg laying during periods of increased Ca<sup>2+</sup> transient frequency in the form of burst firing (Fig. 6B) (Collins *et al.*, 2016; Ravi *et al.*, 2018a). We observed a significant increase in HSN Ca<sup>2+</sup> transient frequency when VC synaptic transmission was blocked compared with nontransgenic control animals (Fig. 6C). WT animals spent ~11% of their time exhibiting high-frequency burst activity in the HSN neurons, whereas transgenic animals expressing TeTx in the VC neurons spent ~21% of their time exhibiting HSN burst firing activity (Fig. 6D). These results are consistent with the interpretation that VC neurotransmission is inhibitory toward the HSNs, such as proposed in previous studies (Bany *et al.*, 2003; Zhang *et al.*, 2008)." |
| **J** | Choi *et al.* 2021 | "VGLUTs are members of a family of anion transporters that move diverse solutes, including inorganic phosphate, acidic sugars, negatively charged amino acids, and phosphorylated adenosine nucleotides<sup>33</sup>. As a member of the SLC17 family of transporters, VST-1 is likely an anion transporter and there are different ways an anion transporter in the synaptic vesicle membrane could limit glutamate uptake. ...we used synaptopHluorin to measure vesicular pH in wild-type and *vst-1* BAG neurons. Measurements of total and surface-accessible pHluorin (Fig. 3g) allow computation of vesicular pH<sub>42</sub>...Importantly, we found that loss of VST-1 caused a measurable increase in vesicular pH (Fig. 3h), consistent with a model in which VST-1 supports anion influx into synaptic vesicles. We also measured vesicular pH in BAG neurons lacking EAT-4/VGLUT (Fig. 3h). Unlike loss of VST-1, loss of EAT-4/VGLUT did not cause a measurable change in vesicular pH. The effect of VST-1 mutation on vesicular pH provides additional evidence that VST-1 functions in the synaptic vesicle membrane. These data are also consistent with a model in which VST-1 is an anion transporter that competes with EAT-4/VGLUT for the electrochemical gradient required for glutamate uptake into synaptic vesicles. However, some SLC17 family transporters can cotransport cations such as Na<sup>+</sup> and H<sup>+</sup> <sup>33</sup>, and we cannot rule out the possibility that cation efflux (rather than anion influx) contributes to the effect of VST-1 on vesicular pH." |
| **K** | Choi *et al.* 2021 | "We further tested whether the effects of *vst-1* mutation on RIA activation by BAGs require GLR-1 glutamate receptors, as predicted by our model. In mutants lacking GLR-1, there was no clear effect of *vst-1* mutation (Fig. 6d, e), indicating that the increased activation of RIAs observed in *vst-1* mutants requires signaling through GLR-1." |

## Results

### *Ce*N-CAM: GO-CAM provides a framework to model neurobiological statements

As a first step in converting information from the scientific literature to a causal model using the GO-CAM framework, we created semantic triples to represent author statements (Supplementary Tables 1 and 2). As an example, a statement by Banerjee *et al*. describing the results of an optogenetic experiment that activates membrane depolarization in uv1 neurons shows that the uv1 cells control the duration of egg deposition during egg-laying behavior. We created a semantic triples to represent this finding: [*membrane depolarization* (GO:0051899)] *occurs in* [*uv1* (WBbt:0006791)] *part of* [*negative regulation of egg deposition* (GO: proposed)] *part of* [*egg-laying behavior* (GO:0018991)] (Supplementary Table 1, local identifier EL12). In creating triples for 123 egg-laying and 98 CO_2_ avoidance author statements, we found that the set of relations used in the GO-CAM data model were sufficient to model all author statements in our dataset. However, we required new classes in several other ontologies (the Gene Ontology (GO), the Evidence & Conclusion Ontology (ECO), and the Environmental Conditions, Treatments & Exposures Ontology (ECTO) (Chan *et al*. 2022)) to describe some statements in both datasets (25/123 statements in the egg-laying dataset, and 84/99 in CO_2_ avoidance) (Table 3). These results show that author statements describing *C. elegans* neurobiology can be faithfully captured using the framework of the GO-CAM data model.

We also found it necessary to re-evaluate some existing definitions and classifications of biological processes under the GO class *behavior* (GO:0007610). For example since the primary term *oviposition* is a subclass of *reproductive behavior* (GO:0019098) in GO and oviposition can be used to describe both the entire behavior of egg laying and to describe the actual deposition of an egg onto a substrate, we requested to switch the primary label of *oviposition* (GO:0046662) with the GO synonym *egg-laying behavior*. We also requested a refinement of the definition of *egg-laying behavior* to ‘A reproductive behavior that results in the deposition of eggs (either fertilized or not) upon a surface or into a medium such as water*’.* In addition, we created a new term *egg deposition* (GO:0160027), defined as ‘The multicellular organismal reproductive process that results in the movement of an egg from within an organism into the external environment*’.* In this way, the mechanical process of *egg deposition* is clearly distinguished from *egg-laying behavior*, which includes its regulation by the nervous system. We requested new terms for the positive and negative regulation of egg deposition, defined as nervous system processes. In addition, we proposed definitions for new classes required to describe CO_2_ avoidance, including *carbon dioxide avoidance behavior* and its parent *behavioral response to carbon dioxide* (Table 3).

Many statements describe findings from genetic perturbations, implicating specific pathways, whereas others, such as cell ablation, leave open a variety of genetic mechanisms by which a phenotype is manifested. Here, we describe the use of different relations and processes to refine models according to the range of conclusions available in each case.

### Statement Category: Linking Neurons, Cellular and Molecular Processes, and Behaviors

Fully elucidating functional neural circuits requires an understanding of the cells (e.g. neurons and muscles) involved in the behavior, the molecular basis of the behavior (e.g. the relevant gene products and their activities), and the coordinated relationships among them to affect the behavior. As with all biological processes, however, the full understanding of a neural circuit and a behavior is produced from individual, granular observations that, together and over time, combine to complete the picture. Leading up to a complete understanding, we need to also have the ability to represent the current state of knowledge at the organismal, anatomical, cellular and molecular level. Thus, in our first category of statements, we aimed to capture atomized statements that link cells and genes to cellular and molecular level processes and those processes to a specific behavior.

A traditional experiment for linking neurons to behavior is to ablate a neuron of interest and observe behavioral effects, an experiment that gives us information at the cellular level (Chalfie *et al*. 1985). When an ablation results in a behavioral change, it is interpreted that one or more processes (either in series or in parallel) occurring in that cell has a causal effect on the behavior (Table 4A). Since cell ablation disrupts unknown cellular processes, we chose to model this result using the high level GO biological process term *cellular process* (GO:0009987), and the *occurs in* (BFO:0000066) relation to contextualize the cellular process with respect to the ablated neuron. We then used the children of the broader causal relation *causally upstream of or within* (RO:0002418) (or preferably a *positive* (RO:0004047) or *negative* (RO:0004046) effect child term) to tie the *cellular process* to a *nervous system process* (GO:0050877). In the example shown in Figure 2A (corresponding to the statement in Table 4A), this *nervous system process* corresponds to the Biological Process term *positive regulation of egg deposition.* We used the *part of* relation in cases where more specific perturbations were made (e.g. neuronal activation or inhibition, genetic knockouts and rescues), allowing an assertion about the composition of the processes involved.

**Figure 2.**
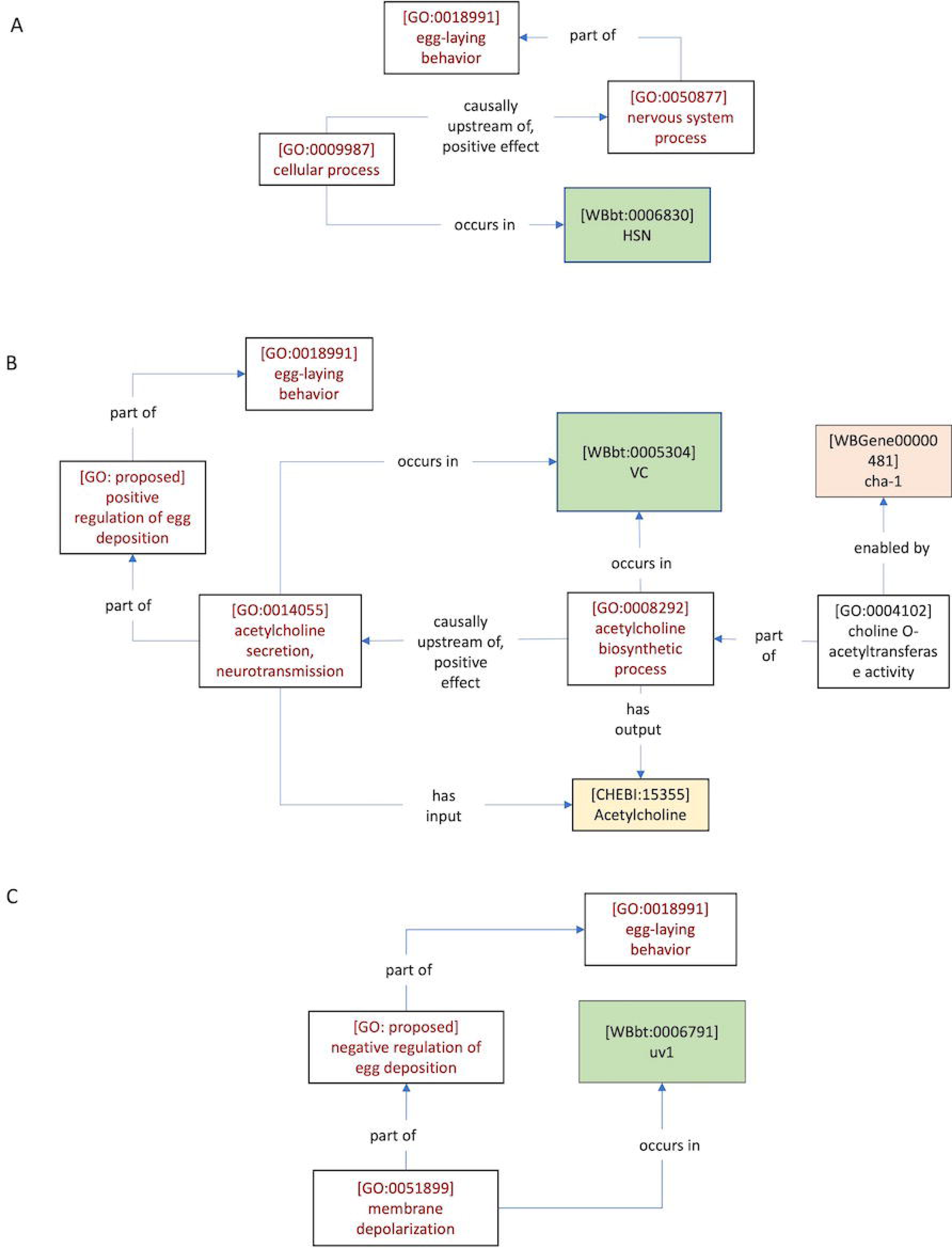
CeN-CAM annotations link cells to behaviors. CeN-CAM annotations link cells to behaviors. Cells such as HSN are represented with unique identifiers in the C. elegans Gross Anatomy Ontology. A) Cell ablation phenotypes can be modelled using the generic GO cellular process class to reflect the non-molecular nature of the experiment, and causally upstream of or within relations, allowing for the most inclusive description of the relationship between cellular process and nervous system process terms. Example drawn from Waggoner et al. (1998) (Table 4A, this manuscript). B) A CeN-CAM model describing the role of the acetylcholine biosynthetic process in the VC neuron in the regulation of egg-laying (Bany et al. 2003) (Table 4B, this manuscript). Because the acetylcholine biosynthetic process can proceed independently of electrical activity, it is modelled as causally upstream of, positive effect; acetylcholine secretion, neurotransmission. C) Optogenetic activation of uv1 leads to a decrease in egg-laying (Banerjee et al. 2017) (Table 4C, this manuscript). This membrane depolarization process is modelled as part of the negative regulation of egg deposition, a nervous system process.

For an illustrative example of this distinction, it is useful to consider experiments from our collection that generated insights by deletion and cell-specific rescue of genes involved in neurotransmitter biosynthesis. We reasoned that since the biosynthesis can proceed even while the neuron is at rest (i.e. independent of the induction of behavior), it should not be considered *part of* the asserted *nervous system process*, but *causally upstream of, positive effect* (RO:0002304) (Fig. 2B, Table 4B) to a secretion process that is *part of* the *nervous system process* (on the assumption that this secretion depends on the depolarization of the neuron).

A more recent experimental technology for discerning the effect of neurons on behavior is optogenetic activation. In these experiments, a specific neuron is activated by opening the light-sensitive Channelrhodopsin ion channel, transgenically expressed in specific neurons of interest (Guo *et al*. 2009). We modelled these results similarly to cell ablation, except that in this case, we were able to say that the *membrane depolarization* that *occurs in* a specific cell is *part of* the *nervous system process* (in this case, the *negative regulation of egg deposition)* that regulates *egg-laying behavior* (Fig. 2C, Table 4C).

### Statement Category: Inputs to Neural Activity & Behavior

A second category of experiment provides insight into the molecular basis of behavior or neural activity induced by an environmental or internal stimulus. In this type of study, a behavior or neural activity that is typically induced by some environmental or experimental (i.e. pharmacological) condition is eliminated under the same conditions when a gene is inactivated. The gene activity is often tied to a cell via rescue of a behavioral mutant phenotype by cell-specific expression of the wild-type allele in the loss-of-function background.

In these cases, we can tie the rescue gene functions to cells, (e.g. in Figure 3A, [*G protein-coupled serotonin receptor signaling pathway* (GO:0098664)] *occurs in* [*VM* (WBbt:0006917)]), and to implied GO biological process terms via *part of* (e.g. in Figure 3B [*intracellular receptor signalling pathway* (GO:0030522)] *part of* [*positive regulation of negative chemotaxis* (GO:0050924)]. In contrast to the case of cell ablation, where unknown cellular processes are disrupted, these more specific biological or cellular process terms can in turn be assigned as *part of* the *nervous system process*. Additional ontology terms and relations can be used to further specify processes or functions. For example, the Chemicals of Biological Interest ontology (ChEBI) contains neurotransmitter classes (e.g. *serotonin* (CHEBI:28790)), as well as environmental chemicals (e.g. *carbon dioxide* (CHEBI:16526) which may be linked to GO receptor activities or other GO molecular functions via *has small molecule activator* (RO:0012001) (Fig. 3A. Table 4D, Fig. 3B, Table 4E).

**Figure 3.**
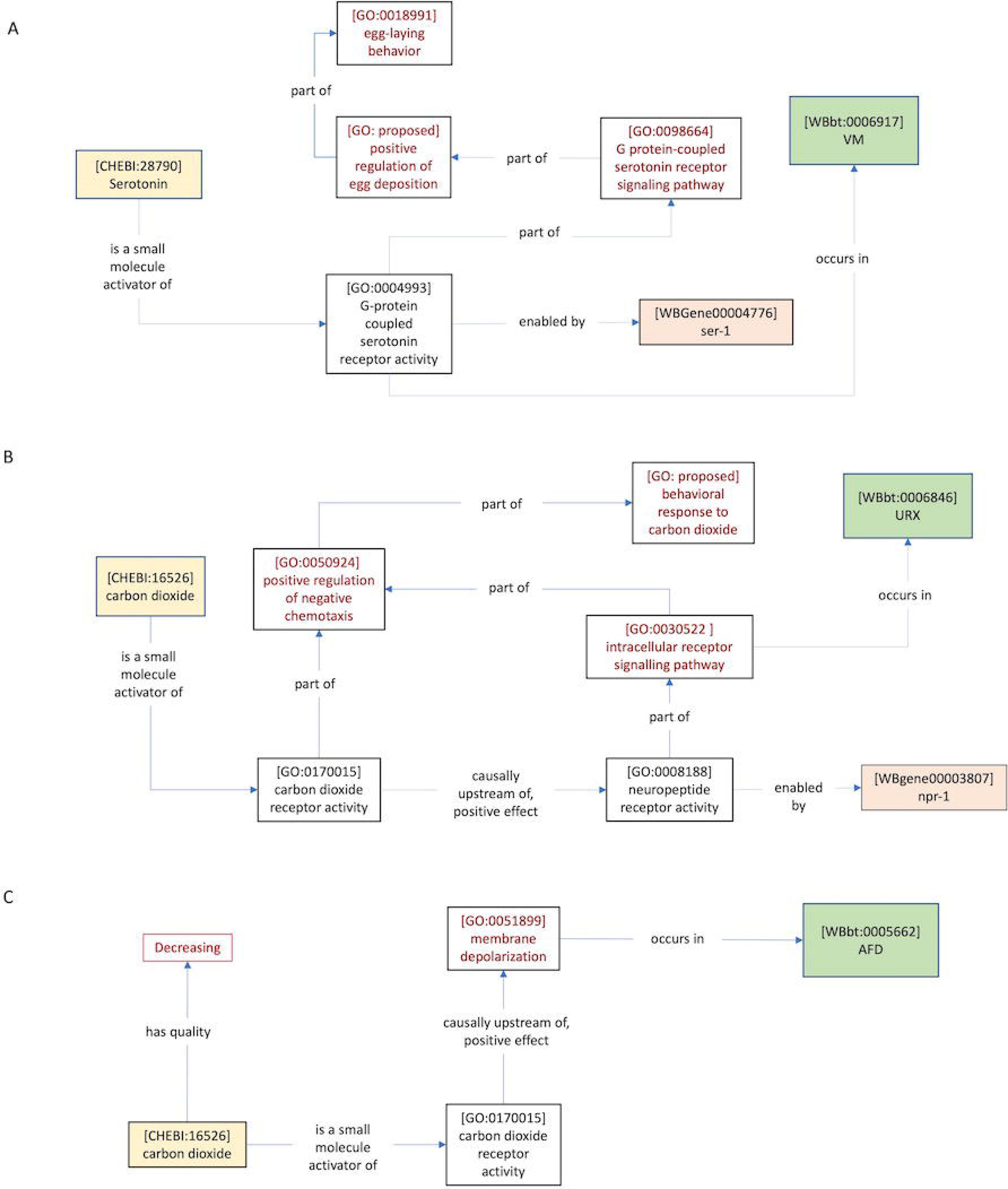
CeN-CAM models of inputs to neurons and behavior. Cell-specific genetic rescue of a behavioral response to pharmacological treatment or environmental stimuli produces models linking genes, GO molecular functions, GO biological processes, and cells in the C. elegans Gross Anatomy Ontology. A) Carnell et al. (2005) (Table 4D, this manuscript) found that VM- specific expression of ser-1 could rescue serotonin-dependent egg-laying behavior, suggesting that ser-1 is required in VM neurons to induce egg-laying in response to serotonin. The G-protein coupled serotonin receptor activity is part of the positive regulation of egg deposition, because the part of relation is transitive (i.e. there is no need for an additional part of relation connecting these nodes). B) Carillo et al. (2013) (Table 4E, this manuscript) found that npr-1 expression in neuronal subsets that include URX is sufficient to rescue the behavioral response to carbon dioxide. The activity of some CO_2_ receptor is implied, leading to the addition of a placeholder term without an enabling gene, indicating an important piece of missing information. This activity can be included in the CO_2_ sensing circuit by asserting that it is part of the positive regulation of chemotaxis, along with the npr-1-dependent signaling pathway. C) AFD neuron responds to CO_2_ removal (Bretscher et al. 2011) (Table 4F, this manuscript). Currently, there are no terms within appropriate ontologies to describe temporal features of chemical or physical inputs (e.g. ‘decreasing’). The required definitions are suggested in this paper.

**Figure 4.**
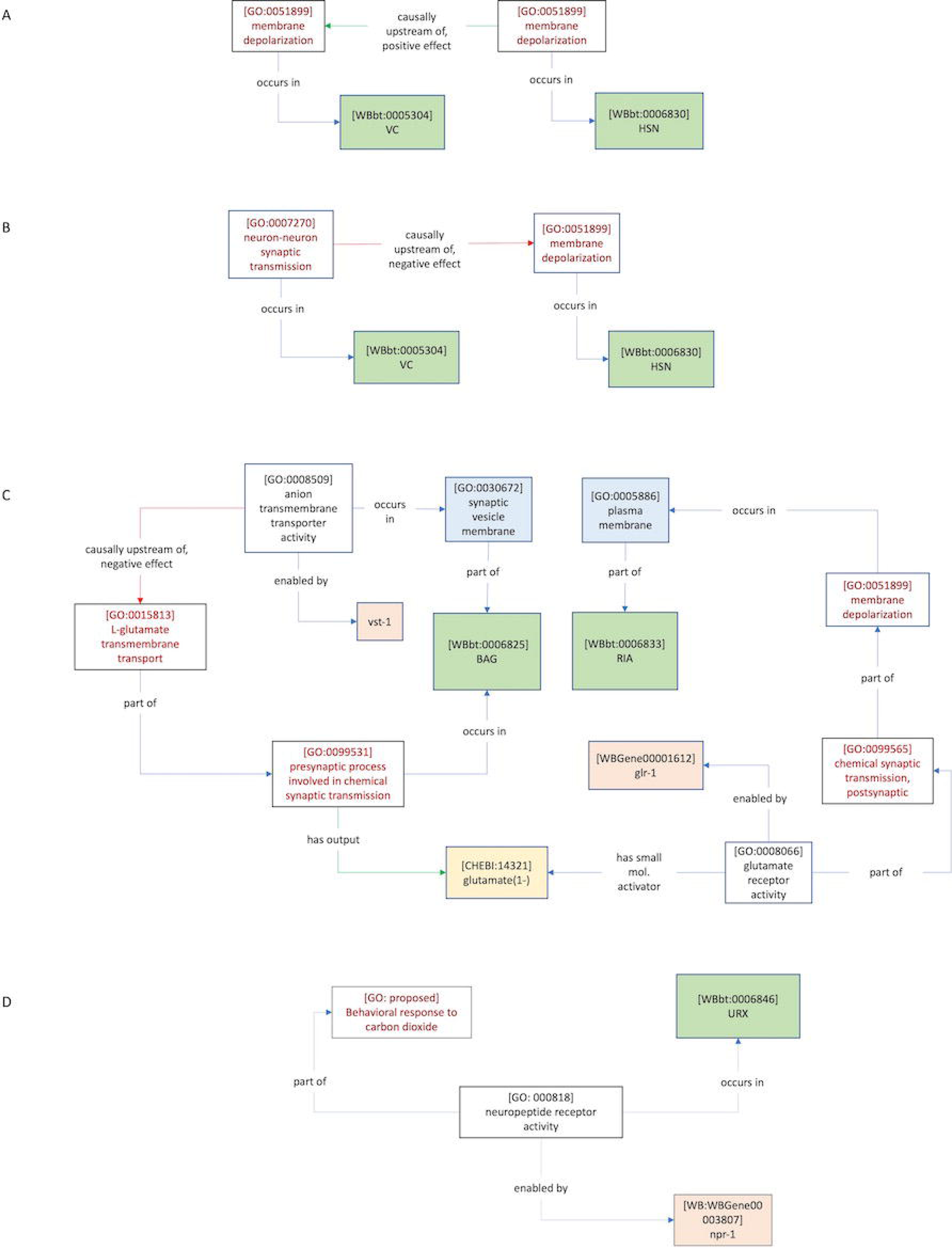
CeN-CAM models of neuron-to-neuron functional connectivity A) Optogenetic activation of HSN neuron causes membrane depolarization in HSN (Collins et al. 2016) (Table 4G, this manuscript). B) Inhibiting neuron-neuron synaptic transmission in VC causes increased activity (membrane depolarization) in HSN, suggesting inhibition of HSN dependent on synaptic transmission from VC (Kopchock et al. 2021) (Table 4H, this manuscript). C) Model of the mechanisms involved in RIA activation based on data from Choi et al. (2021) (Table 4J-4K). Blue boxes represent a GO cellular component term. D) An alternative, more basic model of the same statement modelled in Figure 3C (Table 4F). Figure 3C represents the preferable method

In some cases, the response to a stimulus is measured in a neuron without knowledge of the receptor molecule. For instance, AFD neurons respond to removal of CO_2_, but the experiment does not identify the receptor molecule (Bretscher *et al*. 2011) (Fig 3C) (Table 4F). Because the receptor molecule is unknown, a rescue experiment cannot localise the receptor activity to a cell, meaning that the response may depend on receptor activity in another neuron. This is indicated by the absence of a relationship between the receptor activity and a neuron (similarly, a gene knockout experiment that disrupts neural activity without cell-specific rescue would tie only the membrane depolarization GO term to the neuron). These examples also demonstrate the use of a *nervous system process* term as an intermediate between the *cellular process* terms and the *behavior* terms. For instance, in our model of the role of *npr-1* in the carbon dioxide sensing circuit, a CO_2_ receptor activity is implied, but not tied to a gene or cell (Fig. 3B). However, the *nervous system process* term (*positive regulation of negative chemotaxis* (GO:0050924) provides a natural point of integration by which the receptor activity (and by implication, the cell in which it acts) can be included as part of the same neural circuit. A representative GO-CAM model can be found here^2^.

### Statement Category: Neuron-to-Neuron Functional Connectivity

An additional type of information necessary for fully modelling neural circuits and behaviors is the functional link between neurons. We were able to model statements describing functional connectivity between neurons. For example, an optogenetic experiment in which one neuron is depolarized by a light stimulus and electrical currents are recorded in another neuron may show how a membrane depolarization process occurring in the upstream neuron results in a subsequent membrane depolarization process in the downstream neuron. To capture this relationship, we can connect two *membrane depolarization* (GO:0051899) processes to one another with the *causally upstream of, positive effect* relation (Fig. 4A, Table 4G).

GO also contains classes sufficient to indicate that the transmission occurs through a synapse, when this is explicitly tested by authors. For instance, Kopchock *et al*. (2021) showed a synapse-dependent inhibitory connection between HSN and VC, using tetanus toxin to perturb synaptic transmission. This could be modelled using the GO term *chemical synaptic transmission* (GO:0007268) or one of its children, and the *causally upstream of, negative effect* (RO:0002305) relation to describe the inhibition (Fig. 4B, Table 4H). A similar representation would be appropriate for an experiment describing increase or loss of activity from a recorded neuron in mutants defective for synaptic transmission via mutation of *unc-13* (encodes Munc13), which is required for synaptic vessel exocytosis (Richmond *et al*. 1999). In contrast, mutation of *unc-31* (encodes CAPS), which disrupts dense-core vesicle exocytosis, is required for extra-synaptic transmission (Speese *et al*. 2007)^3^. GO does not have an explicit term for extra-synaptic signaling, or neuropeptide ligand activity. We include an example representation for an extra-synaptic peptidergic connection between two neurons (Supplementary Figure 2C), and provide a definition for the required new GO classes (Table 3). Finally, we include an example that illustrates how *Ce*N-CAM models can represent sub-cellular phenomena involved in neuron-to-neuron functional connectivity in molecular detail (Fig. 4C) (Table 4J- 4K). This model compiles findings from Choi *et al*. (2021), who use the connection between RIA and BAG neurons to investigate mechanisms by which neurotransmitters are loaded into synaptic vesicles.

### Generic Data Models for Statement Categories

In modeling author statements, we found it possible to construct models with varying levels of detail, e.g. cell types, gene products, etc.. For instance, Figure 4D represents a ‘minimal model’ of the same statement described in Figure 3B, representing the rescue of CO_2_ avoidance by expression of the *npr-1* gene in URX. We sought to provide a set of standards for the ideal model of a given category of experimental finding. In our view, a satisfying model will have a structure that corresponds to the conceptual framework of the field (here, the causal flow from inputs to circuits to behavior), and will explicitly illustrate missing knowledge. By modelling the biology that results from different categories of experimental studies, we were able to produce such generic data models for every category (Supplementary Figures 1, 2). In these models, the availability of GO terms and RO relations is constrained by parentage, i.e. only the generic term in the model or one of its children should be used. Importantly, the models are intended to be flexible, i.e. editable using the Noctua GO-CAM modelling software (Thomas *et al*., 2019). In particular, high-level *cellular process* and *nervous system process* terms can be attached to as many GO molecular functions and genes as required to represent the biology. These generic models could accommodate results from both the egg-laying and CO_2_ circuits, suggesting that they may be more broadly applicable to *C. elegans* neurobiology. These models can serve as useful starting points for researchers or biocurators to generate representations of the experimental results, with minimal prerequisite knowledge of the underlying data model.

### GO-CAM can model neural circuits

Systems neuroscience seeks to understand the causal relationships between neural circuits, the behaviors they control, and the inputs that stimulate these circuits, in molecular detail. Having established that a wide variety of author statements describing neurobiological knowledge can be represented in semantic triples, and describing the required GO classes, we generated a model that captures some of the causal relationships within a single circuit. This graph represents interactions between four of the cells that influence egg-laying behavior, from a limited subset of statements in our collection (Fig. 5).

**Figure 5.**
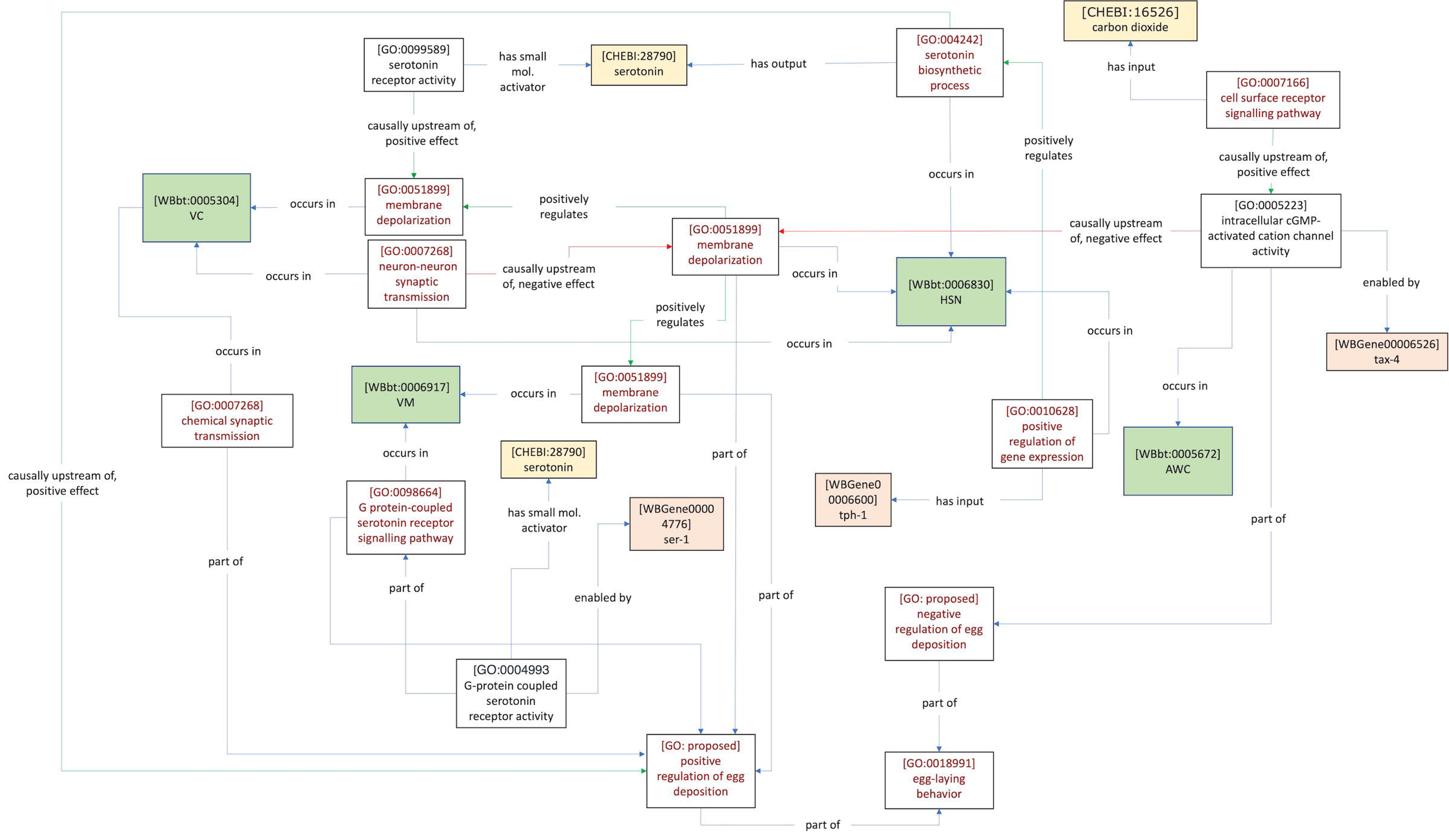
CeN-CAM representation of several cells in the egg-laying circuit and their interactions. Drawn from several statements in Supplementary Table 1 (Carnell et al. 2005; Collins et al. 2016; Fenk & de Bono 2015; Kim et al. 2001; Kopchock et al. 2021). Compared with models of individual author statements, more specific relations can be used here, given the biological context (for instance positively regulates, rather than causally upstream of, positive effect).

Though this diagram does not contain all cells, or all known connections that contribute to egg-laying, it illustrates several useful features of using the GO-CAM framework to model this biology. For instance, the influence of AWC in the circuit is connected to the rescue of HSN inhibition through AWC-specific expression of *tax-4* (Fenk and de Bono 2015). This presumably involves chemical output from AWC that depends on its electrical activity; however, the author statement does not assert this specifically. Similarly, the serotonin synthesized in HSN is likely to be causally involved in the activation of VC, via activity-dependent release into the synapse connecting these two neurons, but this has not been demonstrated directly – only that exogenous serotonin can substitute for the absence of HSN, where there is evidence for *tph-1-* dependent serotonin biosynthesis (Zhang *et al*. 2008). Finally, we used two nodes to represent serotonin, because it allows the possibility that the HSN-VC serotonergic connection may be synaptic, while the HSN- VM connection is extra-synaptic. Thus, *Ce*N-CAM models can represent causal flow within anatomical networks in molecular detail, at the level of what is known, supported by statements in the published literature, and as a result, also indicate what knowledge is missing.

In addition, we show that it is possible to use more informative relations in the context of a model that integrates various findings from the egg-laying literature, compared to those used to model individual author statements. In the case of representing author statements, our models were restricted to the use of information contained in those statements. Here, in the larger *Ce*N-CAM model, we are able to use relations that reflect an overall interpretation of the biology, such as *positively regulates* (RO:0002213) (a child of *causally upstream of, positive effect*) to describe interactions between processes in different neurons.

### GO-CAM can model simple circuit phenomena

Many studies of neural circuits investigate the mechanistic basis for information processing capabilities in the brain, such as the integration of inputs from multiple sensory modalities, and changes in behavior that depend on memory of past experience. We extended our modelling efforts to represent some of these findings, primarily from our CO_2_ avoidance behavior dataset.

### Context-Dependence & Multisensory Integration

An important function of nervous systems in any organism is the ability to execute behavioral responses in a context-dependent manner. This requires integrating multiple kinds of environmental information, ‘computing’ on that information and eliciting an appropriate response. This integration may commonly be performed either by individual neurons responsive to multiple inputs, or by small circuits of three or more neurons, e.g. single interneurons that integrate input from multiple sensory neurons (Ghosh *et al*. 2017). Capturing this type of integration requires relations that imply the necessity of multiple conditions toward a single response, sometimes referred to as AND logic.

We found a relevant example in our CO_2_ avoidance dataset. In one study, *tax-2*-dependent rescue of CO_2_ avoidance was found to depend on the presence of food (Bretscher *et al*. 2011) (Table 5A). We considered whether any of the GO-CAM relations can be interpreted as conveying necessity, in particular the relation *part of.* When considering processes such as those represented in a model, if one process is part of another process, then the latter process necessarily has the former process as a part (or subprocess), meaning that in these contexts *part of* and *has part* (BFO:0000051) are inverse relations (Smith *et al*. 2005). Figures 6A-6C show how the necessity for AND logic might be modelled. The cell-specific rescue of CO_2_ avoidance via *tax-2* expression in BAG neurons, along with the inferred CO_2_ receptor activity, constitutes one ‘branch’ of the model. A second ‘branch’ represents the involvement of food, via an inferred *signal transduction* (GO:000716) process. These two branches converge on a proposed GO term s*ignal integration process* via *part of* relations, capturing their joint necessity. We chose to include a new GO biological process for their integration (rather than having them converge on positive regulation of CO_2_ avoidance) in order to represent that the mechanism enabling the AND logic should be asserted. This representation leaves open many possible biological models for the mechanism by which the asserted integration might occur (for example, one in which food and CO_2_ are sensed by distinct sensory neurons, and integrated in a third interneuron), while capturing AND logic.

**Figure 6.**
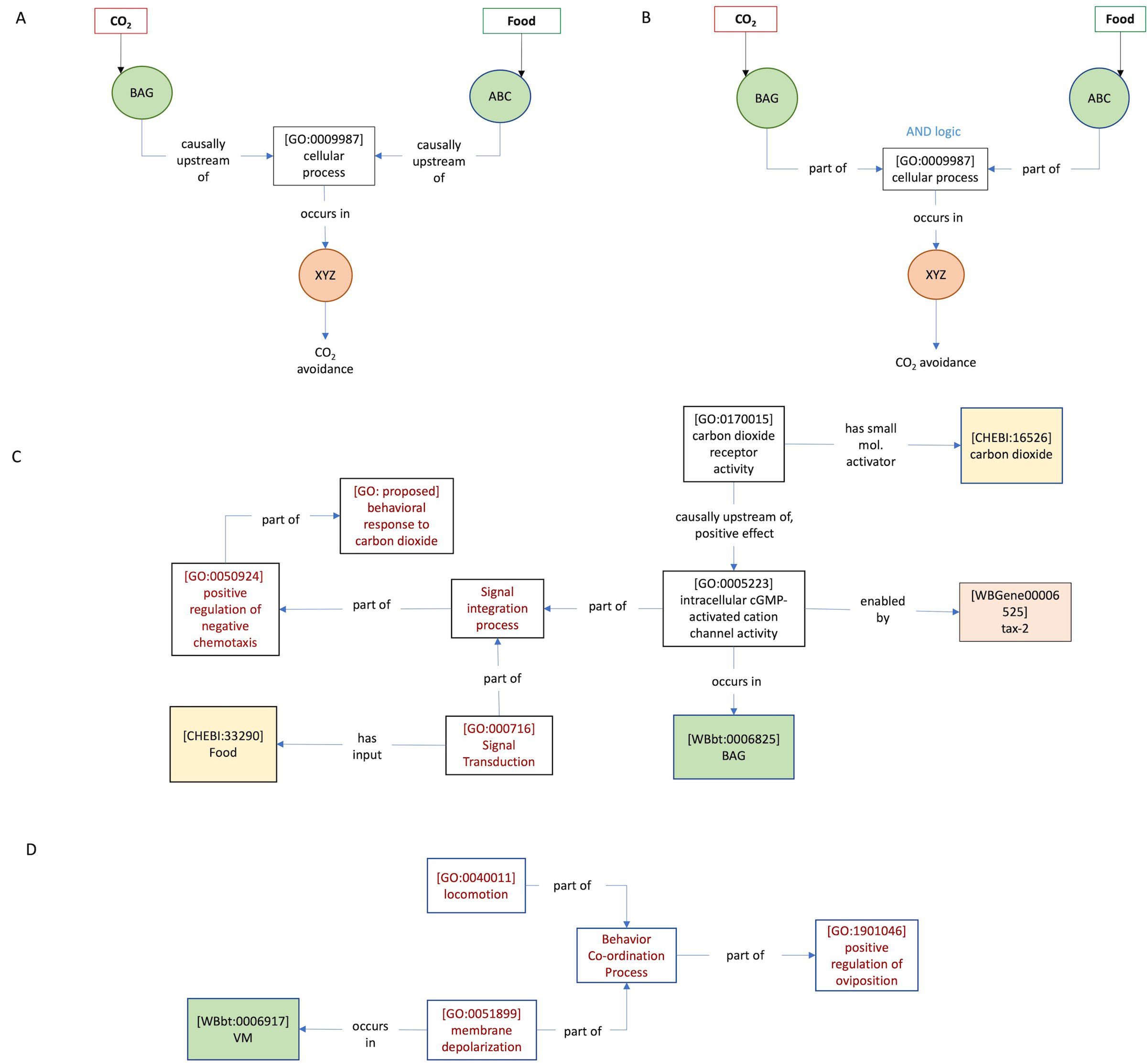
Modelling signal integration with the part of relation. Bretscher et al. (2011) (Table 5A, this manuscript) found that restoring tax-2 expression to BAG neurons rescued CO_2_ avoidance on food, but not off food, suggesting that tax-2-dependent avoidance behavior requires food input. This implies an AND-gated interaction to integrate food and CO_2_ signals. (A) One of the relations in GO-CAM, causally upstream of does not capture the necessity of each input, whereas (B) part of does imply necessity, as required to capture the AND logic involved in sensory integration. (C) GO-CAM representation of the author statement listed in Table 2A (this manuscript). (D) This relation may also be useful for modelling co-ordination of behaviors. Kopchock et al. (2021) (Table 5B, this manuscript) found that optogenetic activation of the vulval muscles was insufficient to induce an egg-laying event; instead co-ordination of VM activation with a particular phase in the body bend during locomotion was required.

**Table 5:**
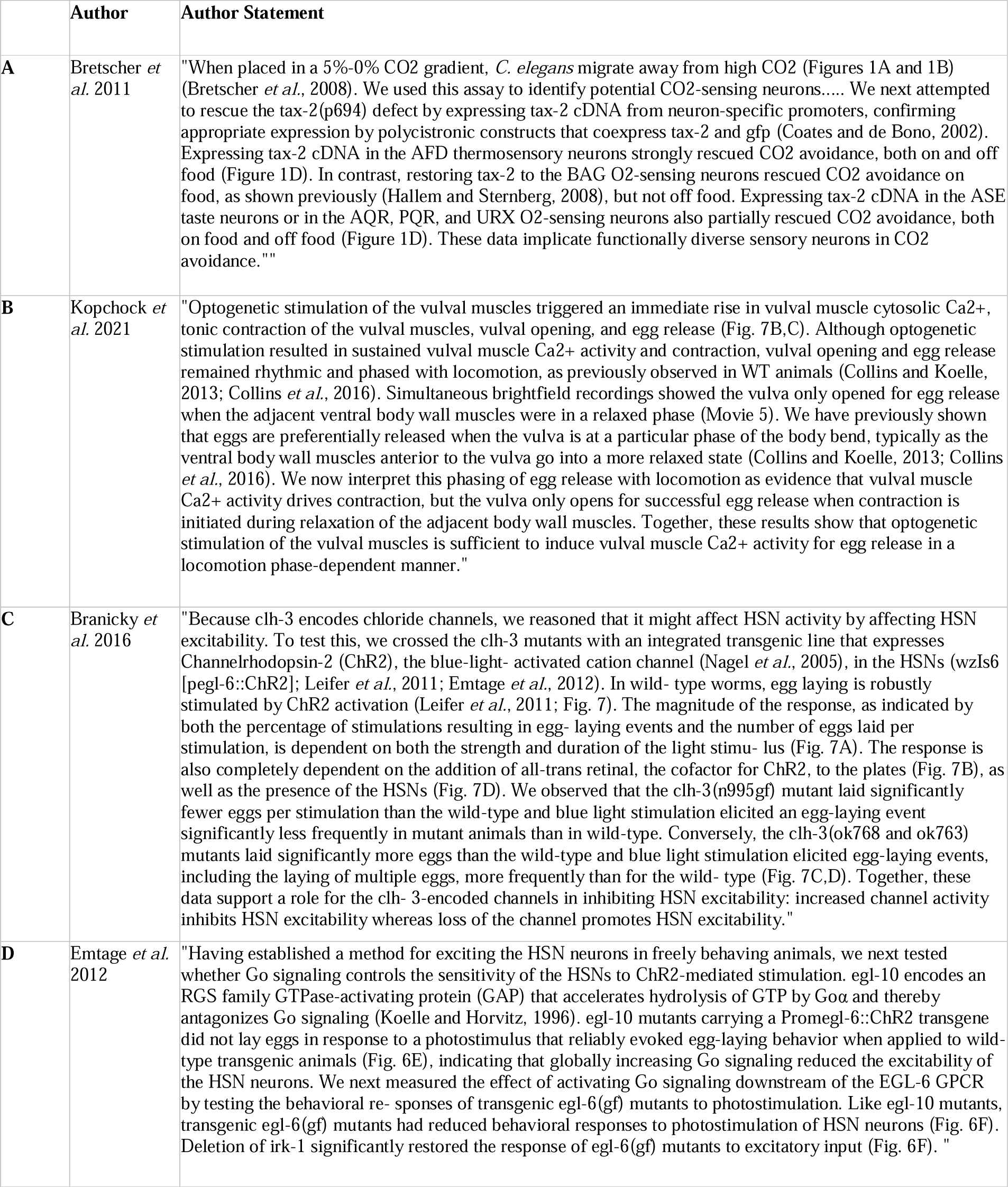
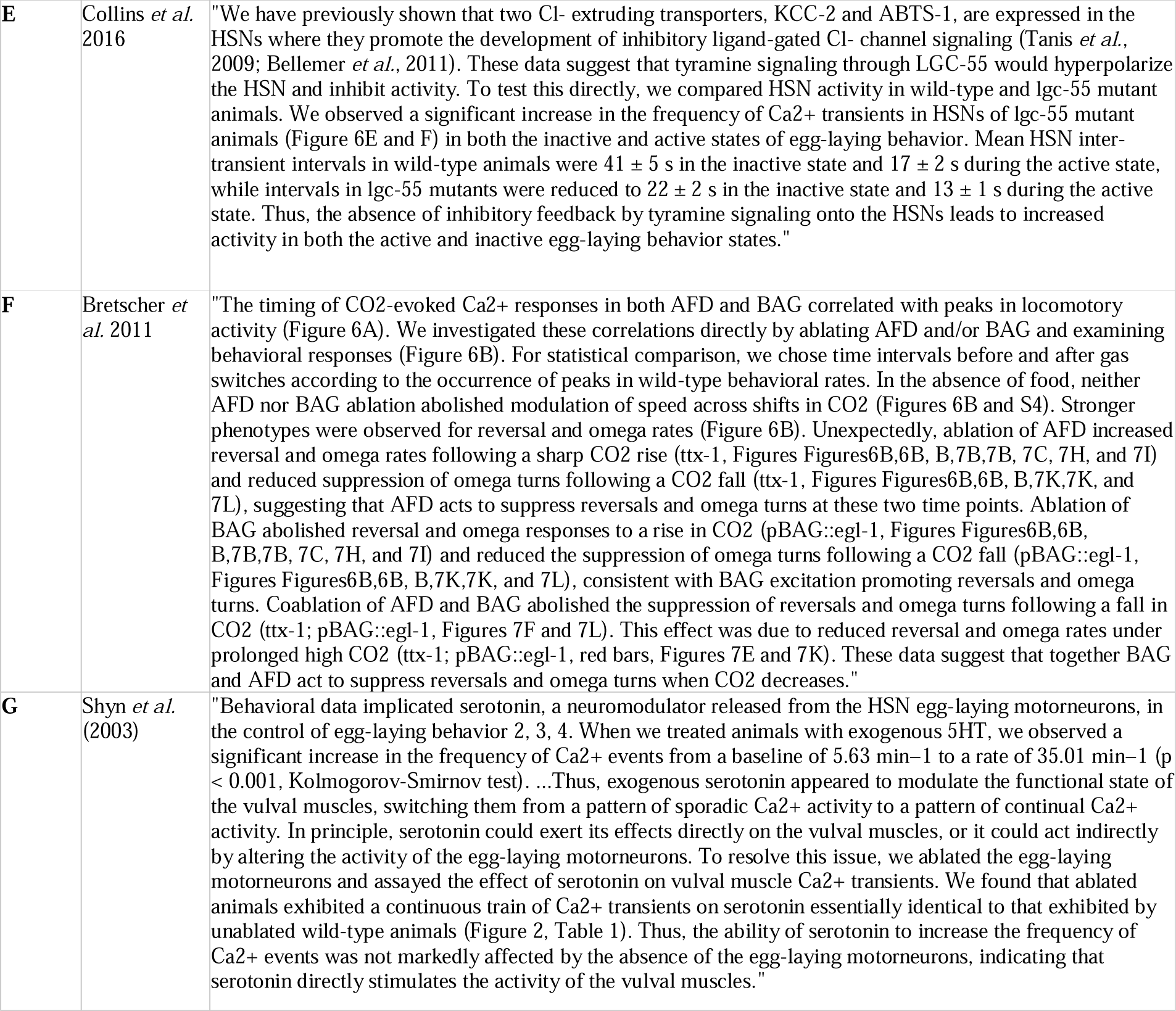
Author Statements Collection B.

|  | Author | Author Statement |
| --- | --- | --- |
| **A** | Bretscher *et al.* 2011 | "When placed in a 5%-0% CO<sub>2</sub> gradient, *C. elegans* migrate away from high CO<sub>2</sub> (Figures 1A and 1B) (Bretscher *et al.*, 2008). We used this assay to identify potential CO<sub>2</sub>-sensing neurons..... We next attempted to rescue the tax-2(p694) defect by expressing tax-2 cDNA from neuron-specific promoters, confirming appropriate expression by polycistronic constructs that coexpress tax-2 and gfp (Coates and de Bono, 2002). Expressing tax-2 cDNA in the AFD thermosensory neurons strongly rescued CO<sub>2</sub> avoidance, both on and off food (Figure 1D). In contrast, restoring tax-2 to the BAG O<sub>2</sub>-sensing neurons rescued CO<sub>2</sub> avoidance on food, as shown previously (Hallem and Sternberg, 2008), but not off food. Expressing tax-2 cDNA in the ASE taste neurons or in the AQR, PQR, and URX O<sub>2</sub>-sensing neurons also partially rescued CO<sub>2</sub> avoidance, both on food and off food (Figure 1D). These data implicate functionally diverse sensory neurons in CO<sub>2</sub> avoidance."" |
| **B** | Kopchick *et al.* 2021 | "Optogenetic stimulation of the vulval muscles triggered an immediate rise in vulval muscle cytosolic Ca<sup>2+</sup>, tonic contraction of the vulval muscles, vulval opening, and egg release (Fig. 7B,C). Although optogenetic stimulation resulted in sustained vulval muscle Ca<sup>2+</sup> activity and contraction, vulval opening and egg release remained rhythmic and phased with locomotion, as previously observed in WT animals (Collins and Koelle, 2013; Collins *et al.*, 2016). Simultaneous brightfield recordings showed the vulva only opened for egg release when the adjacent ventral body wall muscles were in a relaxed phase (Movie 5). We have previously shown that eggs are preferentially released when the vulva is at a particular phase of the body bend, typically as the ventral body wall muscles anterior to the vulva go into a more relaxed state (Collins and Koelle, 2013; Collins *et al.*, 2016). We now interpret this phasing of egg release with locomotion as evidence that vulval muscle Ca<sup>2+</sup> activity drives contraction, but the vulva only opens for successful egg release when contraction is initiated during relaxation of the adjacent body wall muscles. Together, these results show that optogenetic stimulation of the vulval muscles is sufficient to induce vulval muscle Ca<sup>2+</sup> activity for egg release in a locomotion phase-dependent manner." |
| **C** | Branicky *et al.* 2016 | "Because *clh-3* encodes chloride channels, we reasoned that it might affect HSN activity by affecting HSN excitability. To test this, we crossed the *clh-3* mutants with an integrated transgenic line that expresses Channelrhodopsin-2 (ChR2), the blue-light- activated cation channel (Nagel *et al.*, 2005), in the HSNs (wzIs6 [pegl-6::ChR2]; Leifer *et al.*, 2011; Emtage *et al.*, 2012). In wild- type worms, egg laying is robustly stimulated by ChR2 activation (Leifer *et al.*, 2011; Fig. 7). The magnitude of the response, as indicated by both the percentage of stimulations resulting in egg- laying events and the number of eggs laid per stimulation, is dependent on both the strength and duration of the light stimu- lus (Fig. 7A). The response is also completely dependent on the addition of all-trans retinal, the cofactor for ChR2, to the plates (Fig. 7B), as well as the presence of the HSNs (Fig. 7D). We observed that the *clh-3*(n995gf) mutant laid significantly fewer eggs per stimulation than the wild-type and blue light stimulation elicited an egg-laying event significantly less frequently in mutant animals than in wild-type. Conversely, the *clh-3*(ok768 and ok763) mutants laid significantly more eggs than the wild-type and blue light stimulation elicited egg-laying events, including the laying of multiple eggs, more frequently than for the wild- type (Fig. 7C,D). Together, these data support a role for the *clh-3*-encoded channels in inhibiting HSN excitability: increased channel activity inhibits HSN excitability whereas loss of the channel promotes HSN excitability." |
| **D** | Emtage *et al.* 2012 | "Having established a method for exciting the HSN neurons in freely behaving animals, we next tested whether Go signaling controls the sensitivity of the HSNs to ChR2-mediated stimulation. *egl-10* encodes an RGS family GTPase-activating protein (GAP) that accelerates hydrolysis of GTP by *Goa* and thereby antagonizes Go signaling (Koelle and Horvitz, 1996). *egl-10* mutants carrying a Promegl-6::ChR2 transgene did not lay eggs in response to a photostimulus that reliably evoked egg-laying behavior when applied to wild-type transgenic animals (Fig. 6E), indicating that globally increasing Go signaling reduced the excitability of the HSN neurons. We next measured the effect of activating Go signaling downstream of the EGL-6 GPCR by testing the behavioral re- sponses of transgenic *egl-6*(gf) mutants to photostimulation. Like *egl-10* mutants, transgenic *egl-6*(gf) mutants had reduced behavioral responses to photostimulation of HSN neurons (Fig. 6F). Deletion of *irk-1* significantly restored the response of *egl-6*(gf) mutants to excitatory input (Fig. 6F). " |
| E | Collins *et al.* 2016 | "We have previously shown that two Cl<sup>-</sup> extruding transporters, KCC-2 and ABTS-1, are expressed in the HSNs where they promote the development of inhibitory ligand-gated Cl<sup>-</sup> channel signaling (Tanis *et al.*, 2009; Bellemer *et al.*, 2011). These data suggest that tyramine signaling through LGC-55 would hyperpolarize the HSN and inhibit activity. To test this directly, we compared HSN activity in wild-type and lgc-55 mutant animals. We observed a significant increase in the frequency of Ca<sup>2+</sup> transients in HSNs of lgc-55 mutant animals (Figure 6E and F) in both the inactive and active states of egg-laying behavior. Mean HSN inter-transient intervals in wild-type animals were $41 \pm 5$ s in the inactive state and $17 \pm 2$ s during the active state, while intervals in lgc-55 mutants were reduced to $22 \pm 2$ s in the inactive state and $13 \pm 1$ s during the active state. Thus, the absence of inhibitory feedback by tyramine signaling onto the HSNs leads to increased activity in both the active and inactive egg-laying behavior states." |
| F | Bretscher *et al.* 2011 | "The timing of CO<sub>2</sub>-evoked Ca<sup>2+</sup> responses in both AFD and BAG correlated with peaks in locomotory activity (Figure 6A). We investigated these correlations directly by ablating AFD and/or BAG and examining behavioral responses (Figure 6B). For statistical comparison, we chose time intervals before and after gas switches according to the occurrence of peaks in wild-type behavioral rates. In the absence of food, neither AFD nor BAG ablation abolished modulation of speed across shifts in CO<sub>2</sub> (Figures 6B and S4). Stronger phenotypes were observed for reversal and omega rates (Figure 6B). Unexpectedly, ablation of AFD increased reversal and omega rates following a sharp CO<sub>2</sub> rise (ttx-1, Figures Figures6B,6B, B,7B,7B, 7C, 7H, and 7I) and reduced suppression of omega turns following a CO<sub>2</sub> fall (ttx-1, Figures Figures6B,6B, B,7K,7K, and 7L), suggesting that AFD acts to suppress reversals and omega turns at these two time points. Ablation of BAG abolished reversal and omega responses to a rise in CO<sub>2</sub> (pBAG::egl-1, Figures Figures6B,6B, B,7B,7B, 7C, 7H, and 7I) and reduced the suppression of omega turns following a CO<sub>2</sub> fall (pBAG::egl-1, Figures Figures6B,6B, B,7K,7K, and 7L), consistent with BAG excitation promoting reversals and omega turns. Coablation of AFD and BAG abolished the suppression of reversals and omega turns following a fall in CO<sub>2</sub> (ttx-1; pBAG::egl-1, Figures 7F and 7L). This effect was due to reduced reversal and omega rates under prolonged high CO<sub>2</sub> (ttx-1; pBAG::egl-1, red bars, Figures 7E and 7K). These data suggest that together BAG and AFD act to suppress reversals and omega turns when CO<sub>2</sub> decreases." |
| G | Shyn *et al.* (2003) | "Behavioral data implicated serotonin, a neuromodulator released from the HSN egg-laying motoneurons, in the control of egg-laying behavior 2, 3, 4. When we treated animals with exogenous 5HT, we observed a significant increase in the frequency of Ca<sup>2+</sup> events from a baseline of 5.63 min<sup>-1</sup> to a rate of 35.01 min<sup>-1</sup> (p < 0.001, Kolmogorov-Smirnov test). ...Thus, exogenous serotonin appeared to modulate the functional state of the vulval muscles, switching them from a pattern of sporadic Ca<sup>2+</sup> activity to a pattern of continual Ca<sup>2+</sup> activity. In principle, serotonin could exert its effects directly on the vulval muscles, or it could act indirectly by altering the activity of the egg-laying motoneurons. To resolve this issue, we ablated the egg-laying motoneurons and assayed the effect of serotonin on vulval muscle Ca<sup>2+</sup> transients. We found that ablated animals exhibited a continuous train of Ca<sup>2+</sup> transients on serotonin essentially identical to that exhibited by unablated wild-type animals (Figure 2, Table 1). Thus, the ability of serotonin to increase the frequency of Ca<sup>2+</sup> events was not markedly affected by the absence of the egg-laying motoneurons, indicating that serotonin directly stimulates the activity of the vulval muscles." |

### Co-ordination of Neural Activity and Physical Features of Behavior

Our egg-laying dataset contains statements describing a mechanical feature of egg-laying regulation, namely neural activity in the vulval muscles is co-coordinated with the phase of body bending during locomotion (Kopchock *et al*. 2021) (Table 5B). We suggest that *part of* and a new GO Biological Process class *behavior co-ordination process* could be used to model this (Fig. 6D). This representation captures the author’s interpretation that neural activity in VM drives oviposition conditional on features of locomotory behavior.

### Neuromodulation

An important goal of neural modeling is to capture neuromodulatory effects, which may be defined as changes in neuronal excitability or dynamics, due to changes in internal state or external context (Bargmann 2012). We found a small number of entries in our egg-laying dataset that described changes in membrane excitability (e.g. Table 5C, 5D). We chose to model these with the GO term *regulation of resting membrane potential* (GO:0060075), with the view that changing the ability of the cell to maintain its resting potential is the primary mechanism for regulating neuronal excitability. However, it may be more appropriate to use a parent GO class that can model changes in excitability, rather than implying any mechanism. For instance, one might imagine induced changes in receptor expression that could alter excitability or responsiveness, without changing the resting membrane potential (Shine *et al*. 2021). The term *regulation of membrane depolarization* (GO:0003245) and its children may be more appropriate when the mechanism is not known.

### Extending Existing Ontology Classes for Modelling Neurobiology

We found that the use of existing ontologies provided the correct classes for building our models of neural circuitry. However, in some cases we found that additional classes would be useful for a complete and accurate description of the type of biology we are modeling. These proposed additional classes would be added to GO, the Evidence and Conclusion Ontology (ECO) and the Environmental Conditions, Treatments and Exposures Ontology (ECTO) and are listed in Table 3. Evidence supported by four types of experiments, chemical inhibition of neurons via histamine chloride (Pokala *et al*. 2014), inhibition of synaptic transmission (Sweeney *et al*. 1995), mechanical perturbation, and long-term exposure experiments require additional classes in ECO. The categories below describe the biological phenomena that require new GO Biological Process terms, GO Molecular Function terms and ECTO terms to model. Inclusion of these new classes would enrich the kinds of queries that could be supported by *Ce*N-CAM (for instance, we may want a list of all interneurons whose activity is known to be modulated by peptidergic output from ASI neuron). Particularly useful would be the addition of the previously mentioned requirement for GO terms describing extra-synaptic neuropeptide signaling and neuropeptide activity. OBO ontologies are carefully managed, and ontology developers provide processes for the addition of new classes. For instance, we were able to add a GO term for *carbon dioxide receptor activity* (GO:0170015) via the GO GitHub repository by providing the necessary information for its incorporation into the ontology (see https://github.com/geneontology/go-ontology/issues/24994). We discuss other proposed classes below.

### Fine Temporal Dynamics of Neural Activity & Behavior

Many statements described neural activity in fine temporal detail. Experimental treatments are sometimes reported to result in changes to either magnitude, duration and/or frequency of membrane depolarization or hyperpolarization (e.g. Table 5E). In some cases, these phenotypes lead authors to the interpretation that these parameters of a neuron’s behavior are under selection in wild-type organisms, and required to perform the given behavioral task (for instance, changes in the frequency of calcium transients in neurons of the egg-laying circuit are thought to reflect shifts from ‘active’ to ‘inactive’ states of the circuit, reflecting phases of the behavior (Collins *et al*. 2016)). However, the GO class for *membrane depolarization* (GO:0051899) does not distinguish these variations, and related terms such as *positive regulation of membrane depolarization* (GO:1904181) explicitly groups these phenomena together under one term. In the future, it may be useful to have these classes separated into explicit categories for a more comprehensive and informative view of how neural activity is regulated.

Likewise, many assays of egg-laying behavior document its temporal features, dividing it into active and inactive phases, and measuring the effect of various perturbations on their duration and frequency (e.g. Table 4D). In the CO_2_ avoidance literature, a small number of entries described fine details in motor output as a result of neuronal perturbations, such as changes in rates of reversal or frequency of omega turns (Bretscher *et al*. 2011) (Table 5F). We were unable to model these features due to a lack of sufficiently fine-grained GO terms in the Biological Process ontology. However, we note that WormBase has a phenotype ontology to describe behavior in many of the appropriate ways (for example *turning frequency increased* (WBPhenotype:0002313)) (Schindelman *et al*. 2011). Since these are mutant phenotypes and not Biological Processes, these ontology classes are a poor fit for *Ce*N-CAM. Conversion of these phenotype classes into meaningful GO Biological Processes would be helpful to create more fine-grained models of behavior.

### Temporal Features of Environmental Input

In modelling environmental inputs, we found it necessary to model several temporal features. For instance, PATO lacked terms required to model changes in input concentration or intensity over time, as required to model the OFF response to CO_2_ in ADF neurons (Fig 5B). We found that terms in the Environmental

Conditions, Treatments & Exposures Ontology (ECTO) came closer to these requirements (e.g. *exposure to decreased methane* (ECTO:4000005)), but a specific exposure term for many chemicals, such as carbon dioxide, does not exist. We propose and define new classes specifying temporal properties that could be hosted in ECTO (Fig. 7) (Table 3).

**Figure 7.**
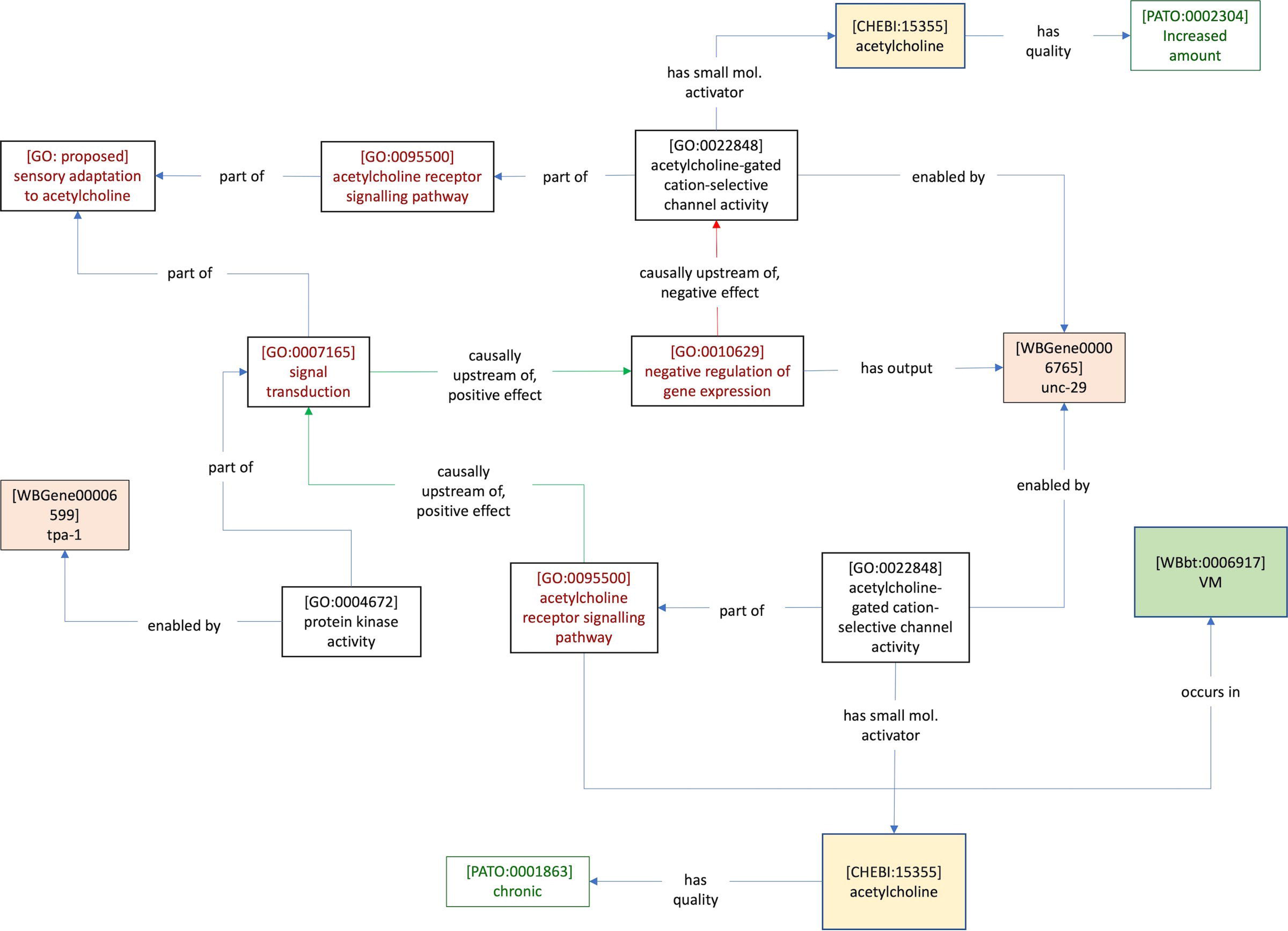
Proposed classes for addition to the Environmental Conditions, Treatments & Exposures Ontology (ECTO) to represent temporal features of environmental inputs. The new classes increasing amount and decreasing amount can be used in combination with the relation has quality (RO:0000086).

## Discussion

Given the size, scope and rapid growth of the biological literature, new methods are required to integrate, represent and interpret accumulating knowledge at varying levels of detail. One method for achieving this is to integrate objects from relevant ontologies in semantic graphs (Thomas et al. 2019; Juanes Cortes et al. 2021). In this work, we demonstrate the applicability of a Gene Ontology based semantic modelling framework, GO-CAM, for representing knowledge of neural circuits in *C. elegans.* By capturing author statements in select papers, we were able to construct simple semantic statements and then link those statements together to begin building causal models of two *C. elegans* behaviors, egg-laying and carbon dioxide avoidance. We found that the existing Relations Ontology (RO) relations used in GO-CAMs are adequate, but new classes are required in several ontologies, including the Gene Ontology (GO), the Evidence & Conclusions Ontology (ECO) and the Experimental Conditions, Treatments & Exposures Ontology (ECTO) to fully represent the statements in our collection.

In general, the GO contains a rich vocabulary for neurobiology, in part due to projects such as SynGO (Koopmans *et al*. 2019), which expanded GO’s representation of synaptic function, and deposited corresponding annotations in the GO repository as GO-CAM models. In addition, the Reactome knowledgebase contains pathways for synaptic transmission, and these have been converted to GO-CAMs (Good et al. 2021). To complement the synaptic transmission part of the ontology, new terms will be required to describe features of extra-synaptic (i.e. peptidergic) connectivity. We also anticipate a more widespread need to model temporal details of sensory neuron input, since chemotactic behaviors typically involve sensing of spatial gradients, experienced by sensory neurons as change over time, resulting in movement towards or away from the odour source. For instance, the sensory neuron AWA adapts to a given concentration of diacetyl, requiring increasing concentration for continued depolarization and associated positive chemotaxis (Larsch *et al*. 2015). Adding these temporal details to the inputs of individual neurons would allow for more expressive representations. In addition, many of the GO Biological Process terms that we propose as additions to the Gene Ontology are the result of describing the processes at the level of an organism or cell and are not derived from attempts to annotate gene function. Such temporal details are often derived from phenotypic measurements resulting from non-genetic perturbation (e.g. cell ablation, pharmacological inputs), in anticipation of the involvement of gene activities in the programmed regulation of these processes. In practice, new GO BP terms based on these observations will likely need genetic evidence before they can be included in the GO; however, we include them here as suggestions, which may guide future proposals as the need arises.

In addition, the models presented here go beyond the minimal requirements for the conversion of author statements into semantic triple format. According to our criteria, a satisfying model should reflect the conceptual framework of the field (in this case, representing causal flow from inputs through circuits to behavior). In this way, the models indicate which knowledge is missing. For instance, Figure 3B depicts the role of *npr-1* in the URX neuron in carbon dioxide avoidance behavior. By including a *nervous system process* term indicating the involvement of neural circuit, it is possible to indicate that a carbon dioxide receptor, whose encoding gene and cellular site of action require identification, are part of the circuit. The data modelling work presented here also provided us with an empirical basis for creating generic models or templates for each of the statement categories described above (Supplementary Figs. 1, 2). In constructing these generic models, we followed structures that reflected the relevant conceptual framework into which particular classes of experimental results should fit. For instance, the full description of a peptidergic connection between neurons should involve the relevant ligand(s), receptor(s), ion channel(s) and encoding genes (Supplementary Fig. 2C). Including the overarching biological process term *neuron-to-neuron signaling by neuropeptide* allows a database to be indexed for these types of connections. In this way, scientists and biocurators can collaborate to generate models with a common understanding of their proper criteria.

We also tried to capture simple ‘computations’ important for nervous system function, and arrived at some modelling principles that are noteworthy. Firstly, when representing the AND logic involved in multisensory integration, it is important to use relations that convey necessity, and have separate causal flows that converge on a single biological process. We note that the proposed GO Biological Process terms (*signal integration process* and *behavior coordination process*) describe an information processing event that could in theory be carried out via any molecular mechanism that satisfies the task. Representing similar kinds of neurobiological knowledge in the GO may require further understanding of the types of molecular mechanisms that typically underlie this type of nervous system process (Ghosh *et al*. 2017).

In this study, we focused on modelling interactions within neural circuits, and their relationship to broad features of behavior, rather than the detailed mechanics of motor programs that they control. In principle, it is possible to link neural activities to the mechanical outputs of neural activity, where both are considered *part of* the organismal behavior under study. In the case of egg-laying, this motor output is simple, involving only the contraction of the vulval muscles. However, CO_2_ avoidance involves a complex series of locomotory processes, each of which is regulated by specific patterns of neural activity (for example, see Bretscher *et al*. 2011). As discussed above, inference of new biological process terms by conversion of the appropriate terms from the *C. elegans* Phenotype Ontology will allow modelling of these features of behavior. These motor outputs could then be modelled as *part of* the organismal behavior *carbon dioxide avoidance behavior* (i.e. they are the targets of the *regulation of chemotaxis* term in the models diagrammed here).

One limitation not previously discussed is that GO-CAM currently has no way of incorporating negative data. In some cases, this prevented documentation of important discoveries from our literature search. For instance, (Shyn *et al*. 2003) found that in the absence of VC neurons and HSN neurons, spontaneous Ca^2+^ transients continued in the vulval muscles, suggesting that these neurons are not necessary for VM activity (Table 5G). These are arguably important omissions from these knowledge graphs.

With these adjustments, this work demonstrates the possibility of creating a machine-readable knowledge base for neurobiology that can return information based on queries. An important part of this resource will be to generate a representation of the *C. elegans* brain that is computable, since the current anatomy ontology does not contain synaptic or gap junction connections between neurons (Lee and Sternberg 2003). Incorporating connectome data that contains the appropriate neuron to neuron relations and property chain algebra (i.e. (Neuron A synapses to Neuron B) and (Neuron B synapses to Neuron C) implies that (Neuron A connects with Neuron C)) will allow queries that include or depend on synaptic connectivity information.

The application and widespread use of this technology depends on the amount of information incorporated into the knowledgebase, much of which at this point is directly dependent on manual input by curators. Given our definition of an author statement as a passage of text following a stereotyped form (hypothesis, observation, interpretation), it is possible to envision how author statements could be identified automatically. We envision a scenario in which machine intelligence could be applied to identify not only author statements, but identify the category of experiment they describe, and the GO terms that correspond to words within them. Using the generic data models described here as templates could help to ensure that machine-generated models are constrained by a desirable structure. With these capabilities, a large volume of the *C. elegans* neural circuit literature could potentially be converted into *Ce*N-CAM models computationally. The author statements that we collected as part of this work will serve as training data to pursue this type of approach.

It is also important for biologists to have usable and intuitive ways of interacting with and analysing synthesized knowledge. One way to achieve this is by representing compiled neurobiological data in an anatomical context. For instance, the Virtual Fly Brain project has used an ontology-based approach to integrate connectivity and single-cell gene expression data, which can be visualised in a 3-dimensional visualization of the brain, using a semantic integration framework (Milyaev et al. 2012; Court et al. 2023). This allows users to run queries to explore gene expression and phenotype data in an anatomic context. We are exploring the possibility of functionally annotating the *C. elegans* connectome in molecular detail using *Ce*N-CAM (Fig. 8). The relevant data are the same as those captured by the statement categories for which we have generated templates, namely causal relationships between inputs to neurons, neurons to behavior, and causal connection between neurons. In addition to populating template data models, the GO terms in the relevant author statements could be used to populate a dataframe of the kind used by visualization software such as Cytoscape (Shannon *et al*. 2003)(Supplementary Figure 3), ideally in an automated manner. This visualization could serve as an intuitive entry point for exploring neural circuit function on a connectome scale, where evidence behind individual elements of the graph could be accessed by linking to the corresponding *Ce*N-CAM models. An anatomical visualization that includes functional and connectivity data would allow predictions to be made about functional relationships between different circuits. For instance, CO_2_ has been shown to inhibit egg-laying (Fenk and de Bono 2015) in an AWC-dependent manner. Representing neurons that respond to CO_2_ along with neurons that control egg-laying in a connectome context (Fig. 9) suggests that ASH is a CO_2_ responsive neuron synaptically linked to HSN. Indeed, ASH was later shown to inhibit both egg-laying and HSN activity (Wen et al. 2020). Functional connectome annotation would also enable various kinds of system-wide analysis of the *C. elegans* brain – a research avenue that has so far been pursued in the absence of functional information (Reigl *et al*. 2004; Alon 2007; Jarrell *et al*. 2012). We also envision the ability to make useful predictions using the underlying semantic models. For instance, the graphs may include causal links between molecular functions and behaviors that result from synthesis of disparate literature, leading to new predictions about how genetic or pharmacological perturbations may affect behavior. Thus, the work described here provides semantically and biologically rigorous foundations for an integrated systems neuroscience resource combining knowledge representation, connectome annotation and associated computational analyses of *C. elegans* nervous system function.

**Figure 8.**
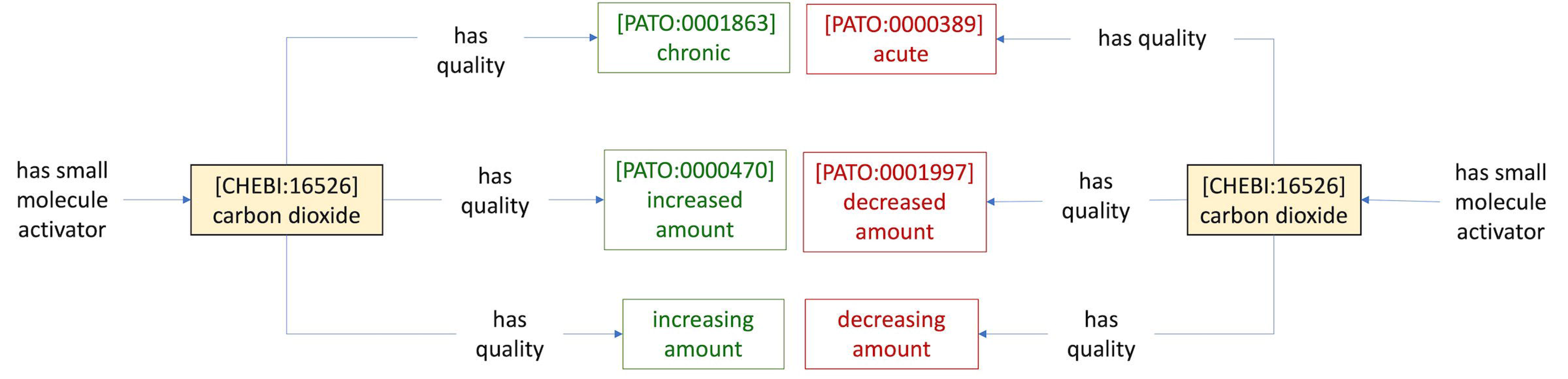
Functional annotation of the C. elegans connectome allows visualization of causal relationships within and among neural circuits. Black or grey arrows indicate synaptic connections from electron microscopy of serial sections. Coloured solid arrows indicate activating (green) or inhibiting (blue) synaptic connections. Coloured dotted arrows indicate activating or inhibiting indirect or extra-synaptic connections. Bold outlines on neurons indicate the sign of the neuron on the specified behavior. Fill colour on neurons indicates the effect of the specified input on neural activity. A) Synaptic connectivity for subset of neurons involved in both CO_2_ avoidance behavior and egg-laying behavior. B) Neurons activated or inhibited by CO_2_ are indicated by fill color, neurons contributing to CO_2_ avoidance behavior indicated by outline color. C) Neurons activated or inhibited by serotonin are indicated by fill color, neurons contributing to egg-laying behavior indicated by outline color. D) Modelling of neural circuits and behavior involves representing causal connections between inputs to neurons, neurons to neurons, and neurons to behavior. These are the same data required for functional annotation of the connectome. The combination of rigorous data modelling integrated with connectome visualization could provide a useful resource for systems neuroscience.

**Figure 9.**
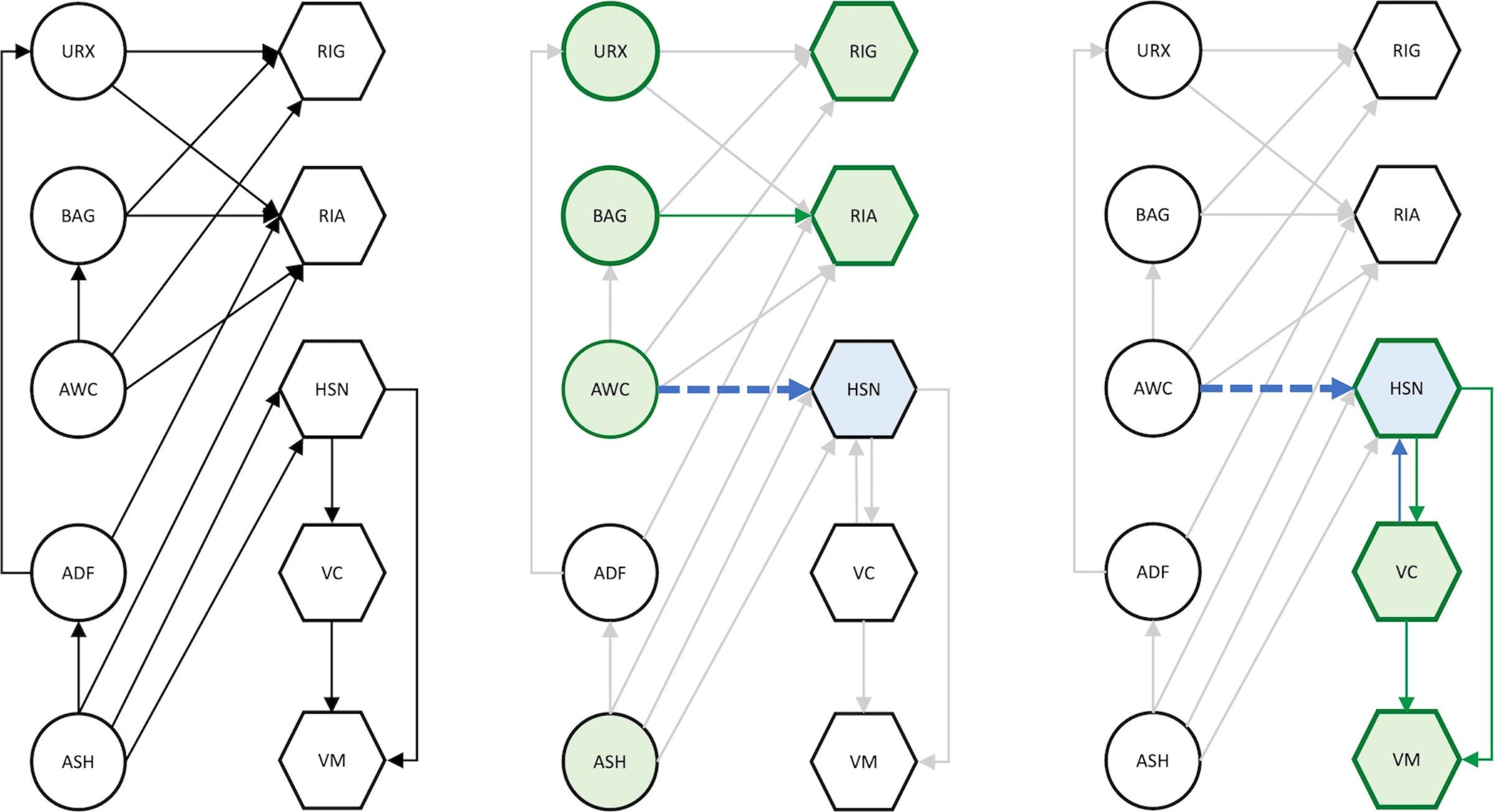

## Declarations

### Abbreviations

GO (Gene Ontology), GOC (Gene Ontology Consortium), GO-CAM (Gene Ontology Causal Activity Modeling), CeN-CAM (*Caenorhabditis elegans* Neural Circuit Causal Activity Modeling), ChEBI (Chemicals of Biological Interest Ontology), Relations Ontology (RO), Evidence & Conclusions Ontology (ECO), Basic Formal Ontology (BFO), Phenotype and Trait Ontology (PATO), Environmental Conditions, Treatments and Exposure Ontology (ECTO).

### Data & Materials Availability

All data generated or analysed during this study are included in this published article (and its supplementary information files).

### Author Contributions

P.W.S & S.J.P. conceived and designed the study, S.J.P, K.V.A & D.P.H. conducted the study. S.J.P, K.V.A, D.P.H & P.W.S wrote and revised the manuscript. S.J.P. prepared all the figures. All authors approved the manuscript.

## Acknowledgements

We thank all members of the Sternberg lab at Caltech for their feedback during the course of the project. We also thank Raymond Lee (WormBase) and members of the Gene Ontology Consortium, as well as Susan Bello (Mouse Genome Informatics) and members of the Unified Phenotype Ontology working group for helpful discussions.

## Footnotes

1 Predicate and relation have the same meaning (predicate is the formal term for describing a triple).

2 http://model.geneontology.org/64e7eefa00000614

3 There is still debate in the literature as to how *unc-13* and *unc-31* may regulate distinct or common processes in synaptic and extra- synaptic transmission (for instance, see Sieburth *et al*. (2007)). In addition, tetanus toxin may disrupt dense core vesicle exocytosis as well as synaptic vesicle exocytosis, as in humans (Hoogstraaten *et al*. 2020). Our modelling here reflects the interpretations of the authors.

