## Supplementary Information for "Semantic Representation of Neural Circuit Knowledge in *Caenorhabditis elegans*"

### Tables

**Supplementary Table 1**
Conversion of author statements to semantic triple format, with statements drawn from the literature on egg-laying behavior. Columns labelled Process # contain nodes representing either GO Biological Processes or GO Molecular Functions. Terms without ID's are novel terms.

**Supplementary Table 2**
Conversion of author statements to semantic triple format, with statements drawn from the literature on carbon dioxide avoidance behavior.

### Figures

**Supplementary Figure 1** Curation templates for experimental results linking neurons to behavior. Each GO term and each relation represents the most generic (highest level) term that is suitable for the model. Authors can populate a model with either these relations, or any of its child terms. High level GO biological process terms can also be expanded to include an arbitrary number of constituent GO molecular functions or GO biological processes. The following labels describe the causal flow described by each model, and the corresponding type of experiment. **A)** Neuron to Behavior (ablation) **B)** Neuron to Behavior (activation/inhibition). **C)** Input to Neuron to Behavior (rescue)

**Supplementary Figure 2** Curation templates for experimental results inputs describing functional connections between neurons, whether **A)** mechanism agnostic (neuron to neuron), **B)** via synapses, or C) extra-synaptic neuropeptide signalling. Terms without ID's are novel terms.

**Supplementary Figure 3** Cytoscape rendering of the same neural circuit as in Figure 9 (this manuscript). Filled cells are responsive to $CO_2$ (green cells are activated, blue cells are inhibited). Outlined cells are involved in egg-laying (green outlined cells promote, and blue outlined cells inhibit egg-laying respectively).
