## Supplementary figures and images for "Semantic Representation of Neural Circuit Knowledge in *Caenorhabditis elegans*"

### Supplementary Figures

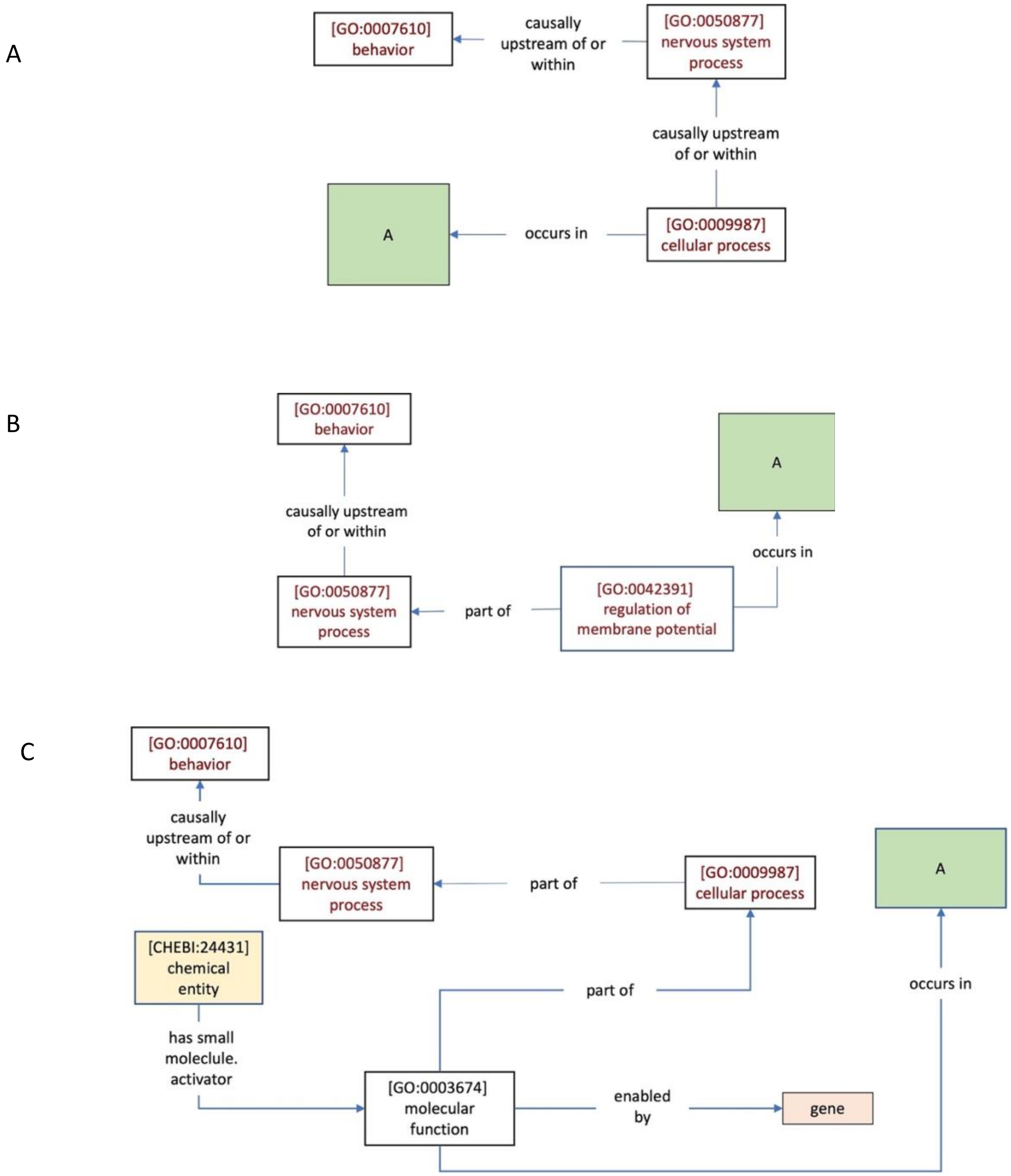


*Supplementary Figure 1*

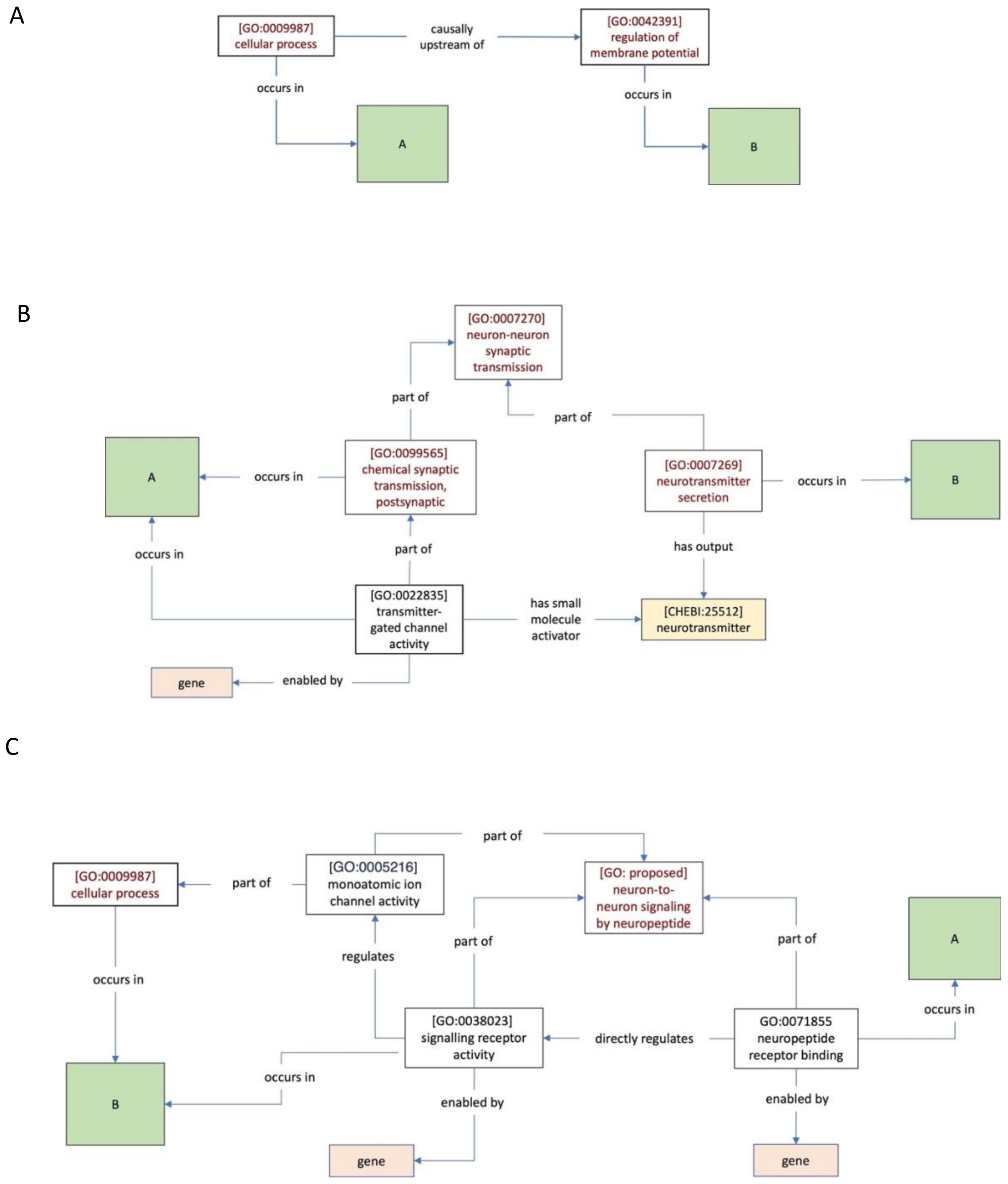


*Supplementary Figure 2*

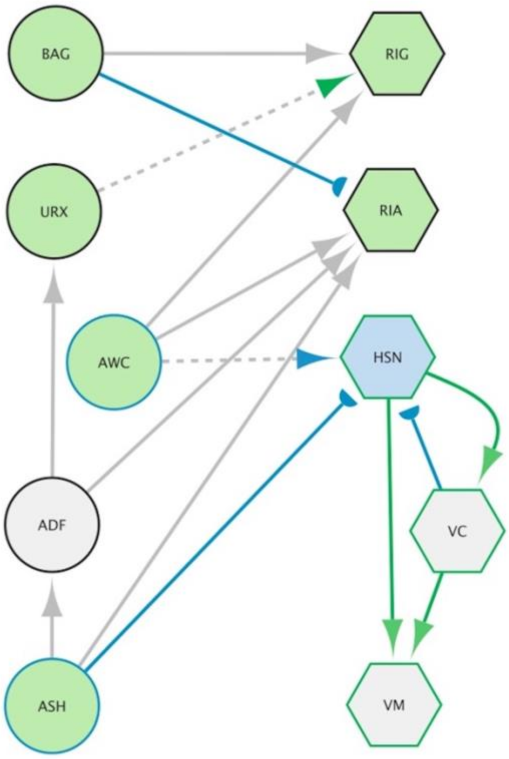


*Supplementary Figure 3*
